# The renal capsule, a vibrant and adaptive cell environment of the kidney in homeostasis and aging

**DOI:** 10.1101/2023.05.11.540033

**Authors:** Ben Korin, Shimrit Avraham, Reuben Moncada, Terence Ho, Mayra Cruz Tleugabulova, Hari Menon, Spyros Darmanis, Yuxin Liang, Zora Modrusan, Cecile Chalouni, Charles Victoria, Linda Rangell, Charles Havnar, Will Ewart, Charles Jones, Jian Jiang, Debra Dunlap, Monika Dohse, Andrew McKay, Joshua D Webster, Steffen Durinck, Andrey S Shaw

## Abstract

The kidney is a complex organ that governs many physiological parameters. It is roughly divided into three parts, the renal pelvis, medulla, and cortex. Covering the cortex is the renal capsule, a serosal tissue that provides protection and forms a barrier for the kidney. Serosal tissues of many organs have been recently shown to play a vital role in homeostasis and disease. Analyses of the cells that reside in these tissues have identified distinct cell types with unique phenotypes. Here, we characterized this niche and found that it is mainly comprised of fibroblasts and macrophages, but also includes other diverse cell types. Characterizing renal capsule-associated macrophages, we found that they consist of a distinct subset (i.e., TLF^+^ macrophages) that is nearly absent in the kidney parenchyma. Injury, disease, and other changes that involve the kidney, affected the cell composition of the renal capsule, indicating its dynamic response to changes within the organ parenchyma. Lastly, we studied age-related changes in the renal capsule and found that aging affected the cell composition and inflammatory phenotype of macrophages, increased CD8 T cells and other lymphocyte counts, and promoted a senescence-associated phenotype in fibroblasts. Taken together, our data illustrate the complexity and heterogeneity of the renal capsule and its underlying changes during aging and disease, improving our understanding of the kidney serosa that may be valuable for novel renal therapies.

## Introduction

Serosal membranes provide structural coverage for many organs and cavities of the body. In each organ, serosal tissues have specific properties that support the normal function of the organ, such as physical support and protection, passage of nutrients, and the secretion of serous fluid to lubricate movement of the underlying organ parenchyma. Recently, serosal tissues of the hepatic capsule^1^, lung pleura^2^, intestine^3^, and brain meninges^4^ have been analyzed. These have been shown to comprise important cellular niches in health and disease.

In homeostasis, cells of serosal tissues are mainly comprised of fibroblasts, mesothelial cells, and macrophages. Fibroblasts play a key role in collagen production, matrix rearrangement, and fibrotic processes, while mesothelial cells cover the outer serosal surfaces and provide a non-adhesive protective barrier of cells that produce the serosal fluid. Within the serosa, macrophages are the predominant immune cell type and present a distinct phenotype compared to circulating or parenchymal macrophages^5^. Macrophages are important for the maintenance of homeostasis, and also take part in injury and repair processes^6, 7^. Thus far, serosal macrophages have been mainly divided into two subsets according to the expression of Lyve-1^low/high^ and MHC-II^high/low^. The absence of Lyve-1^high^MHC-II^low^ macrophages is correlated with increased lung and heart fibrosis, suggesting a role in tissue inflammation^5^. In intraperitoneal bacterial infection, liver serosal macrophages have been shown to mediate neutrophil recruitment to control intrahepatic bacterial dissemination^1^. GATA6^+^ peritoneal resident macrophages have important roles in immune surveillance in the peritoneal cavity, host defense, tissue repair, and peritoneal tumorigenesis^8^. For example, Gata6^+^ resident peritoneal macrophages can promote the growth of liver metastasis^9^. Peritoneal serosal Lyve-1^+^ macrophages have also been shown to affect cancer outcomes as syngeneic epithelial ovarian tumor growth is strongly reduced following *in vivo* ablation of Lyve-1^high^ macrophages^10^. Additionally, in the brain meninges, parenchymal border macrophages (PBMs), that also express high levels of Lyve-1, have gained attention in how they regulate cerebrospinal fluid dynamics during aging and Alzheimer’s disease^11^.

Despite the recent interest in serosal tissues, the cell dynamics and functions of the kidney serosa, the renal capsule, in injury, disease, and aging are less known. The renal capsule is a tough fibrous layer of collagen and elastin that covers and supports the kidney along with the peripheral adipose tissue that surrounds it. The renal capsule has been previously suggested as a stem cell niche in the kidney, possibly participating in the repair of renal injury^12^. During inflammatory and autoimmune conditions of the kidney, subcapsular lymphoid-like structures are present, and potentially promote inflammatory processes in the kidney^13^. Here, we characterize cells of the renal capsule, identifying stromal, myeloid, and lymphoid cells that can adapt and respond to internal changes within the injured, diseased, or aged kidney.

Recently, Dick at el^14^ showed that most tissues have interstitial macrophages that comprise three major subsets: TLF^+^ (TIMD4^+^LYVE1^+^FOLR2^+^), CCR2^+^ (TIMD4^neg^LYVE1^neg^FOLR2^neg^), and MHC-II^high^ (TIMD4^neg^LYVE1^neg^FOLR2^neg^CCR2^neg^), each with unique transcriptional and functional elements, life cycles, and origins. While the TLF^+^ subset self-renews with only a small monocyte contribution, the CCR2^+^ and MHC-II^high^ subsets are mainly monocyte-derived^14^. Using this taxonomy, we found that renal capsule-associated macrophages (RCAMs) are mostly comprised of the TLF^+^ subset. We also found that the dynamics and composition of immune cells in the capsule changes with glomerular injury, indicating a possible crosstalk between the different parts of the kidney.

Since age is a significant risk factor for chronic kidney disease, we also characterized the renal capsule in aged animals and humans. A common characteristic of renal aging and disease is hardening of the renal capsule. We found a substantial increase in certain lymphoid and myeloid cells in the renal capsule of aged mice, and the macrophage transcriptional phenotype became more pro-inflammatory. Single cell RNA sequencing (scRNA-Seq) and flow cytometry of renal capsule cells showed a shift in the balance of macrophage subsets in the renal capsule, with age resulting in decreased TLF^+^ macrophage numbers and increased numbers of CCR2^+^ and MHC-II^high^ subsets. Computational analysis suggested this shift may result from crosstalk between renal capsule lymphoid cells, fibroblasts, and macrophages. Our data suggest the renal capsule as an adaptive and diverse niche within the kidney.

## Results

### Cell composition and phenotype of the renal capsule

To identify the main cell types within the renal capsule of adult mice (C57BL/6J), we analyzed single cell suspensions of the tissue using mass cytometry, flow cytometry, and scRNA-Seq (**Figure 1a, Supplementary Table 1,2**). Using these methods, we identified a variety of immune and non-immune cells: macrophages, monocytes, neutrophils, dendritic cells (DCs), B cells, CD4 and CD8 T cells, γδ T cells, NK cells, fibroblasts, endothelial cells, and epithelial cells (**Figure 1b,c; Supplementary Figure 1a-d**). Macrophages and fibroblasts were most abundant, and we confirmed the presence of these cells in the mouse and human renal capsule by imaging whole-mounted tissue (**Figure 1d**, **Supplementary Figure 1e-g**). Despite the clear difference in cell proportions and composition compared to that found in the murine blood, we validated that the analyzed cells were not contaminated with blood using three approaches. First, we compared the cell composition of renal capsules taken from perfused and non-perfused mice (**Supplementary Figure 2a**). Second, we injected fluorescently labelled anti-CD45 antibody intravenously shortly before harvesting the kidneys, to label cells in the vasculature (**Supplementary Figure 2b**). Third, we used antibodies to the red blood cell marker TER-119 (**Supplementary Figure 2c**). All experiments indicated that there was minimal blood contamination in our renal capsule cell preparations.

**Figure 1:**
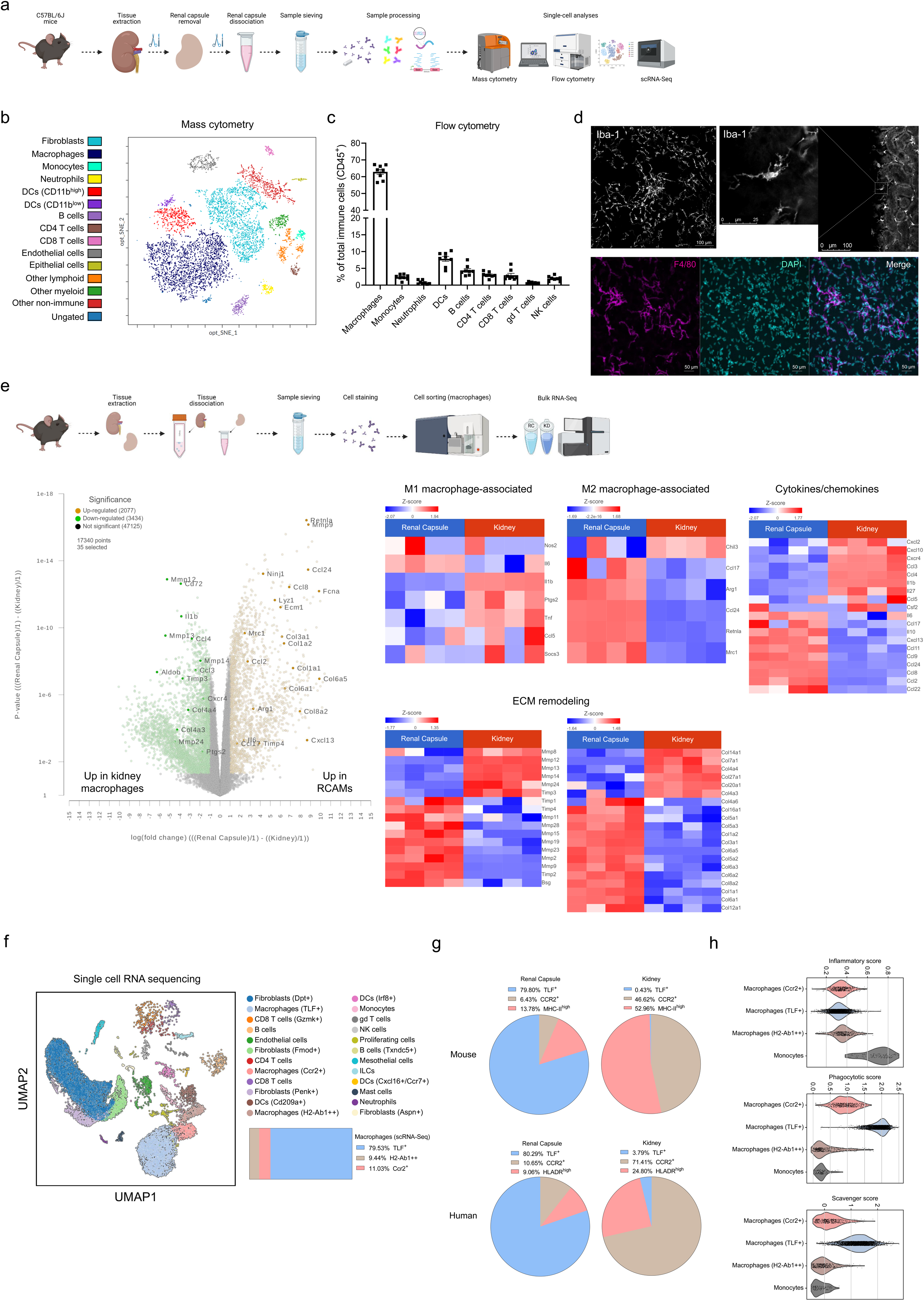
Cell populations of the renal capsule. (**a**) Schematic description of the experimental design for single cell-based analyses. Created with BioRender.com. (**b**) opt-SNE plot of mass cytometry data depicting cell populations in the renal capsule (n=10 pooled mice). (**c**) Flow cytometry data of main immune cell types in the renal capsule (n=8 mice). (**d**) Iba-1 immunofluorescence imaging of macrophages in a whole-mounted mouse renal capsule (top left) and Iba-1 immunofluorescence imaging in tissue-cleared mouse kidneys (top right). F4/80 immunofluorescence imaging in a whole-mounted mouse renal capsule with DAPI (bottom). (**e**) Schematic description of the experimental design for RCAM and kidney macrophages characterization using bulk RNA-Seq. Created with BioRender.com. Volcano plot and heat maps of bulk RNA-Seq data of renal capsule macrophages and kidney macrophages with highlighted selected genes, representative image of two experiments (n=4,4). (**f**) UMAP of scRNA-Seq analysis of renal capsule cells, and horizontal slice chart with the proportions of 3 main interstitial macrophage subsets according to the scRNA-Seq data (2 samples of n=4 pooled mice each). Representative image of two experiments. (**g**) Pie charts with the proportion of 3 main interstitial macrophage subsets in mouse (n=6) and human (25-year-old donor) renal capsules and kidneys, according to flow cytometry analysis. (**h**) Inflammatory and phagocytic path scores of monocytes and macrophages, as previously described.

Since macrophages were the predominant cell population within the renal capsule, we focused on characterizing them in detail. For simplicity, and to maintain consistency with following results, we termed the entire macrophage population in the renal capsule as renal capsule associated macrophages (RCAMs). We first independently sorted RCAMs and kidney macrophages and compared them using bulk RNA-Seq (**Figure 1e**). Transcriptionally, RCAMs were substantially different from kidney macrophages (**Supplementary Figure 3a,b**), presenting a more anti-inflammatory (M2-like) phenotype, while differing in the expression of several cytokine and extracellular matrix (ECM) remodeling genes, such as *Il1b*, *Il10*, *Ccl2*, metalloproteases, and collagens (**Figure 1e**). Unlike peritoneal macrophages, RCAMs did not express *Gata6* (**Supplementary Figure 3c**). Additionally, unbiased high-dimensional cell surface marker analysis (opt-SNE) using mass cytometry highlighted the differences between RCAMs and kidney macrophages, showing, for example, that CD206 and CD169 were higher in RCAMs and CD14 and CD11c were lower compared to kidney macrophages (**Supplementary Figure 4a-c)**.

### TLF^+^ macrophages are the main macrophage subset of the adult murine and human renal capsule

To deepen our characterization of macrophage subsets and other cell types within the renal capsule, we used scRNA-Seq. We validated the presence of fibroblasts (four clusters), CD8 T cells (two clusters), CD4 T cells, B cells (two clusters), endothelial cells, DCs (three clusters), monocytes, γδ T cells, NK cells, neutrophils, and macrophages (three clusters), as well as identified mesothelial cells, ILCs, mast cells, and a cluster of proliferating cells (**Figure 1f**, **Supplementary Figure 5a-d**). Of the three macrophage clusters, we identified macrophage subsets that correspond to the previously described^14^ TLF^+^ (*Timd4^+^Lyve1^+^Folr2^+^*), MHC-II^high^, and CCR2^+^ subsets (**Figure 1f**, **Supplementary Figure 5b**). The proportion of TLF^+^ macrophages in our scRNA-Seq analysis was distinct from that previously described in the kidney (renal capsule: 79.53% (**Figure 1f**), kidney: <1% of macrophages^14^). We confirmed this result by performing flow cytometry of TLF^+^ macrophages from mouse (CD11b^+^CD64^+^Timd4^+^Lyve1^+^Folr2^+^) and human (CD11b^+^CD64^+^Timd4^+^Folr2^+^) renal capsules versus kidneys. This verified that macrophages of the renal capsule are highly enriched for TLF^+^ macrophages compared to the kidney parenchyma (**Figure 1g**). Using a functional macrophage classification system developed by Sanin et al.^15^, monocytes had the highest ‘inflammatory score’ while TLF^+^ macrophages had the highest ‘phagocytic score’ (**Figure 1h**). Using key scavenger-related genes, we also plotted a ‘scavenger score’, which was highest in TLF^+^ macrophages (**Figure 1h**). Given the importance of IL-34 and colony stimulating factor 1 (CSF-1), the ligands of the CSF-1 receptor (CD115) in macrophage maintenance and survival, we examined the expression of *Csf1* and *Il34* in our data and identified that they are mainly expressed by renal capsule fibroblasts, with relatively higher levels of *Csf1* expression (**Supplementary Figure 5d**).

Revisiting our bulk RNA-Seq analysis of RCAMs for TLF^+^ macrophage markers, we confirmed higher expression of TLF-associated genes (as described in^14^) in RCAMs compared with kidney macrophages (**Supplementary Figure 6a**). This was further validated at the protein level by imaging for Lyve-1 and CD206 (**Supplementary Figure 6b**), and by flow cytometry that showed higher expression of multiple TLF^+^ macrophage markers in mouse and human RCAMs, including Lyve-1, CD206 and the scavenger receptor CD163 (**Supplementary Figure 6c**; see **Supplementary Figure 6d** for flow cytometry gating strategy of TLF^+^ macrophages).

### Internal changes within the kidney affect renal capsule immune cell composition

Our results so far have shown that the renal capsule is a complex cell environment that encompasses a variety of immune cell types. Analyzing the immune cell composition of human renal capsules showed the presence of similar cell types as found in mouse, with abundant macrophages (**Figure 2a-c**). These data also suggested that cells in the renal capsule change in response to stresses on the kidney (e.g., in obesity **Figure 2b** or trauma, **Figure 2c**). Therefore, to test this hypothesis, we analyzed the immune cells in the renal capsule using different conditions in mice (**Figure 2d-g**, **Supplementary Figure 7a-j**). We used macrophage MHC-II expression as a functional parameter of antigen presentation (**Supplementary Figure 7a,b,d,f**). First, we queried whether sex influences the immune cell composition of the renal capsule by comparing renal capsules from male and female C57BL/6J mice. Male and female mice demonstrated modest differences in macrophage and B cell percentages (**Figure 2d**, **Supplementary Figure 7a**). Next, to investigate whether major physiological changes within the kidney affect the composition of renal capsule immune cells, we analyzed immune cells following several models that are known to affect kidney function: gestation, obesity, and immune-mediated glomerular injury.

**Figure 2:**
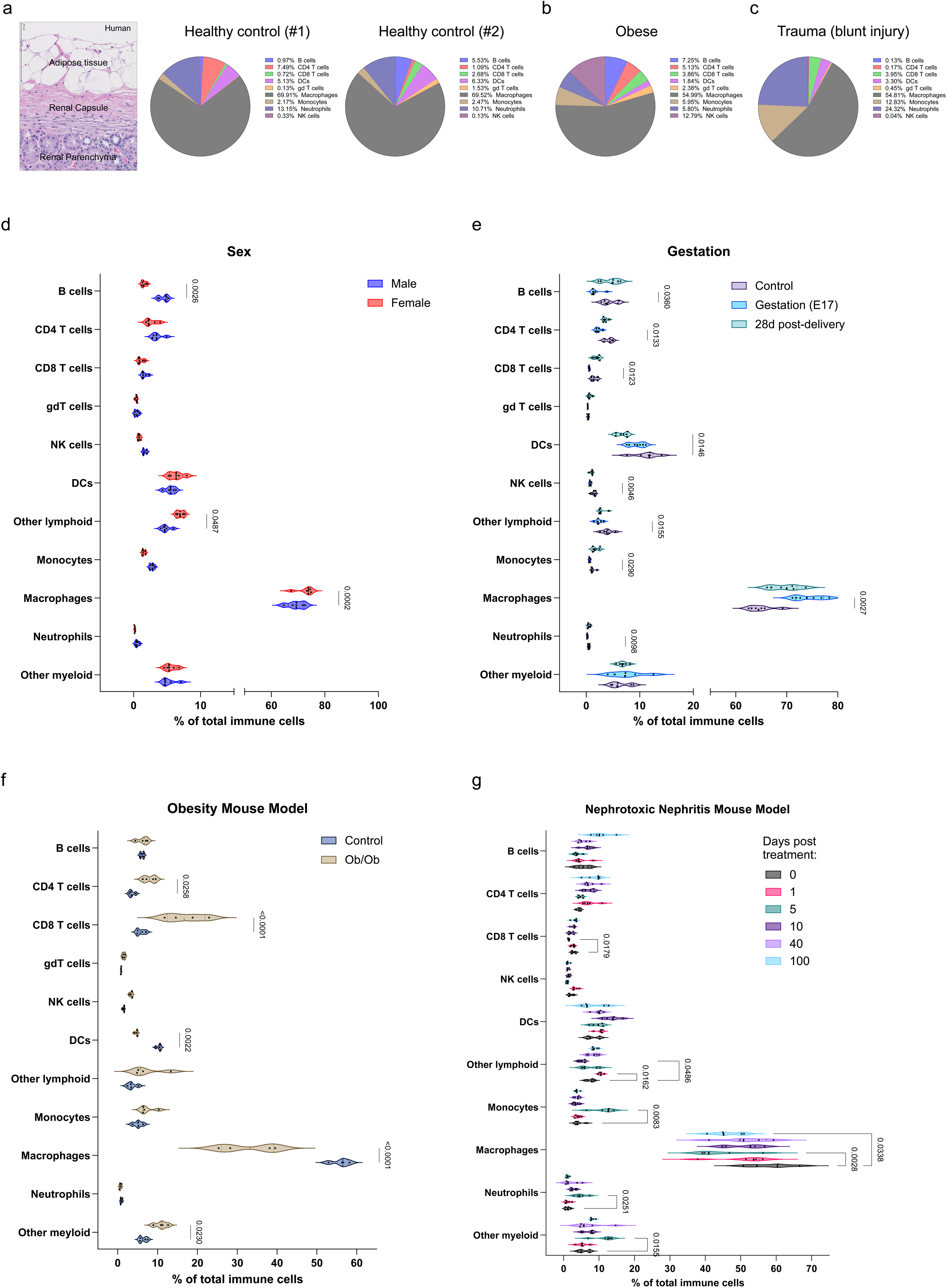
Renal capsule immune cell composition and dynamics in human samples and in different mouse models and conditions. (**a-c**) Representative histology (H&E) image of the outer cortical area in a healthy human kidney, and pie charts with the percentage of main immune cell types in human renal capsules of four donors (left to right): (**a**) left: male 37y BMI=36.1, right: male 25y BMI=23.3, (**b**) male 54y BMI=44.6, (**c**) male 12y BMI=19.1 (cause of death: physical trauma), analyzed by flow cytometry. (**d-g**) Flow cytometry analysis indicating the percentage of main immune cell populations in the renal capsules of: (**d**) 3-month-old male and female C57BL/6J mice (n=5,5, two-way ANOVA). (**e**) Pregnant (E17), 28 days post gestation and control female C57BL/6J mice, 4-6 months of age (n=6,6,6, two-way ANOVA). (**f**) Obese (B6.Cg-Lepob/J) and control mice (1 year-old; n=4,4, two-way ANOVA). (**g**) Nephrotoxic nephritis (NTN) and control C57BL/6J mice, 3-6 months of age (0, 1, 5, 10, 40, and 100 days following nephrotoxic serum (NS) intravenous injection; n=5 per time point, two-way ANOVA).

In human and mouse gestation, kidneys undergo anatomical and hemodynamic changes, to accommodate for increased blood volume, blood pressure regulation, and fetal care^16, 17^. These include increased glomerular filtration rate (GFR) and larger overall organ volume due to increased vascular and interstitial volume^16^. Since kidney volume changes require changes in the renal capsule, we examined the composition of cells in the renal capsule at different stages of gestation. Kidneys of control (non-pregnant), E17, and 28 days post-delivery female mice were harvested, and immune cells in the renal capsule were analyzed. We identified several changes in immune cell proportions, especially when compared to the E17 samples (**Figure 2e**). Renal capsule macrophages were much more abundant at E17 and at 28 days post-delivery compared to controls. Lymphocyte percentages were decreased at E17 and returned to normal post-delivery (**Figure 2e**). However, MHC-II expression in macrophages was lower at E17 and remained so at 28 days post-delivery (**Supplementary Figure 7b**), which may suggest suppressed antigen presentation state of these cells.

To explore the effects of metabolic changes on the renal capsule, we utilized the leptin deficient mouse model of diabetes (ob/ob). This mouse develops insulin resistant diabetes characterized by obesity (**Supplementary Figure 7c**), glucose intolerance, elevated plasma insulin, and hyperglycemia. Renal injury in these mice begins around 10 weeks of age with the onset of proteinuria. Pathological lesions are evident by 20 weeks of age. We analyzed mice at 12 months of age, a time point when they were clearly obese. Here, we identified increased proportions of CD4 and CD8 T cells, accompanied by a decrease in myeloid cells (DCs and macrophages) (**Figure 2f**). The expression of MHC-II was unchanged (**Supplementary Figure 7d**).

Lastly, we used the nephrotoxic serum nephritis (NTN) model of immune complex-mediated glomerular disease^18^. In this model, an anti-serum containing antibodies against the glomerular basement membrane (GBM) proteins (nephrotoxic serum) is injected intravenously. The antibodies bind and accumulate at the glomerular basement membranes initiating an acute immune-mediated injury to the glomerulus with significant proteinuria at day 1 that was largely resolved by day 10-14 (**Supplementary Figure 7e**). Mice were challenged with nephrotoxic serum and renal capsules harvested on day 0, 1, 5, 10, 40 and 100. There were a variety of changes to cell composition that were induced by nephrotoxic serum injury (**Figure 2g**). Most notable was the increased percentage of neutrophils and monocytes at day 5, and the decreased percentage of macrophages on days 5 and 100 post treatment (**Figure 2g**). As the renal inflammation has long resolved, this suggests that the memory of the injury persists in the cells of the capsule.

### Aging increases inflammation in the renal capsule and changes the RCAM phenotype

It is well established that the kidney is highly affected by aging; many kidney diseases are strongly age-associated, especially chronic kidney disease. Aging kidneys are characterized by decreased renal function due to accumulated loss of nephrons, declining glomerular filtration and increased fibrotic scaring^19–21^. To identify aged-related changes within the renal capsule, we extracted kidneys of 6-, 12-, 18-, and 24-month-old mice and scored the level of inflammation, fibrosis, and injury in the renal capsule by pathology. As age increased, we identified higher scores in all histological parameters (**Figure 3a**; for detailed scores see **Supplementary Figure 8a**, **Supplementary Table 4**), which was associated with increased T cell infiltration (**Supplementary Figure 8b**). Analysis of the renal capsule cells by flow cytometry showed increased immune cell numbers in the aged (24-month-old) mouse renal capsule compared to young (3-month-old), especially in CD8 T cells (**Figure 3b**, **Supplementary Figure 8c,d**). These results were consistent with increased RNA expression levels of *Cd3d*, *Cd3g*, *Cd5*, *Cd8a*, *Cd8b1* in samples of whole renal capsules (**Supplementary Figure 8e**). While the total macrophage cell count was similar between young and aged animals, bulk RNA-Seq of sorted young and aged RCAMs showed that the transcriptional phenotype of aged RCAMs was substantially different when compared to the young. For example, aged RCAMs had increased expression of triggering receptor expressed on myeloid cells 2 (*Trem2*), and Apolipoprotein D (*Apod*), both previously associated with a variety of metabolic diseases and chronic inflammation, and vascular cell adhesion molecule 1 (*Vcam1*) which affects endothelial cell interactions (**Figure 3c**, **Supplementary Figure 8f**). Pathway analysis of the top differentially expressed genes identified the induction of vascular, cell migration, communication, and adhesion pathways (**Supplementary Figure 8g**). Additionally, we identified a pro-inflammatory phenotypic shift that was characterized by increased expression of M1-associated genes, decreased expression of M2-associated genes (**Figure 3d**), and elevation of cytokines (e.g., *Cxcl13*, *Il1b*) and cytokine receptors (e.g., *Cxcr4*) genes (**Figure 3e**). Several ECM remodeling-related genes were also higher in aged RCAMs compared with young (**Figure 3f**).

**Figure 3:**
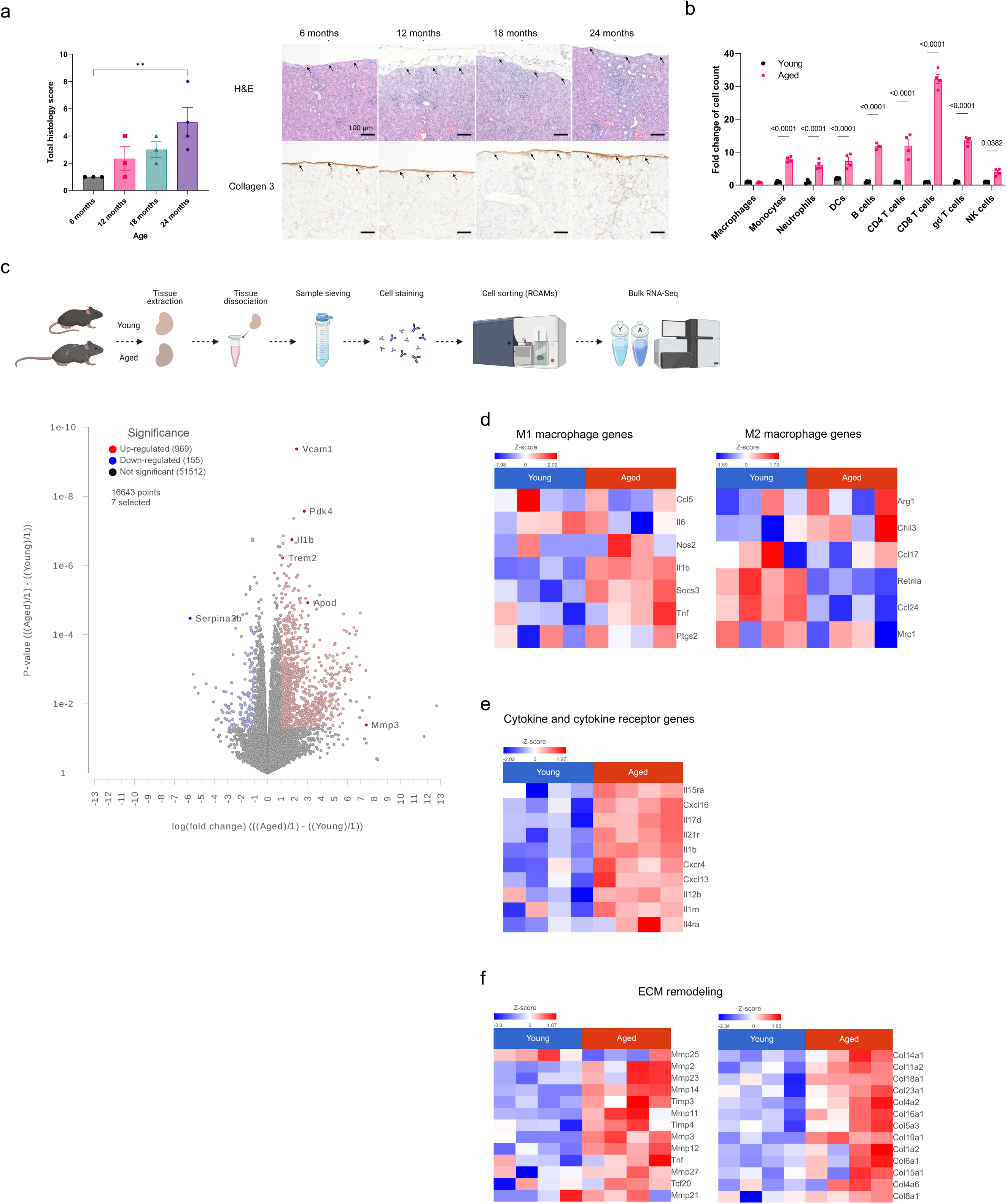
Age-related changes in the renal capsule and its macrophages. (**a**) Renal capsule focused histological analysis of total histology score according to inflammation, tubular atrophy, fibrosis and injury of 6-24 months old mice (n=3,3,3,4, one-way ANOVA, Fisher’s LSD posttest; complete analysis parameters that determined the overall histology score are available in Supplementary Figure 8a), and representative images of H&E and collagen 3 immunolabeling (black arrows point to the renal capsule). (**b**) Flow cytometry analysis of the fold change of cell counts of renal capsule immune cells from young (3-month-old) and aged (24-month-old) mice (n=4,4; each point represents n=4 young and n=3 aged pooled renal capsules; two-way ANOVA). (**c**) Illustration of the experimental design for bulk RNA-Seq of sorted RCAMs of young and aged mice. Created with BioRender.com. Heatmap comparison of (**d**) M1 and M2 hallmark genes, (**e**) matrix metalloprotease genes and extracellular matrix genes, (**f**) main cytokine and cytokine receptors genes.

### Aging reduces TLF^+^ macrophage numbers and compromises their function while promoting senescence-associated CD8 T cells and fibroblasts

To obtain better granularity of the biological processes occurring during ageing, we performed scRNA-Seq of cells from young and aged mouse renal capsules (**Figure 4a**, **Supplementary Figure 9a**). Consistent with the flow cytometry data, the scRNA-Seq analysis of young (3-month-old) versus aged (24-month-old) renal capsules showed changes in lymphoid and myeloid cell proportion and counts, most prominently an increase in the *Gzmk*^+^ CD8 T cell subset (**Figure 4b**, **Supplementary Figure 9b**). Staining for TER-119 suggested little to no blood contamination (**Supplementary Figure 9c**). The percentage and numbers of all fibroblast clusters (*Penk^+^*, *Fmod^+^*, *Dpt^+^*, *Aspn^+^*) were lower in the aged mouse renal capsule when compared to young (**Figure 4b**, **Supplementary Figure 9b**). This could potentially be attributed to hardening of the capsule and decreased accessibility of fibroblasts to digestion.

**Figure 4:**
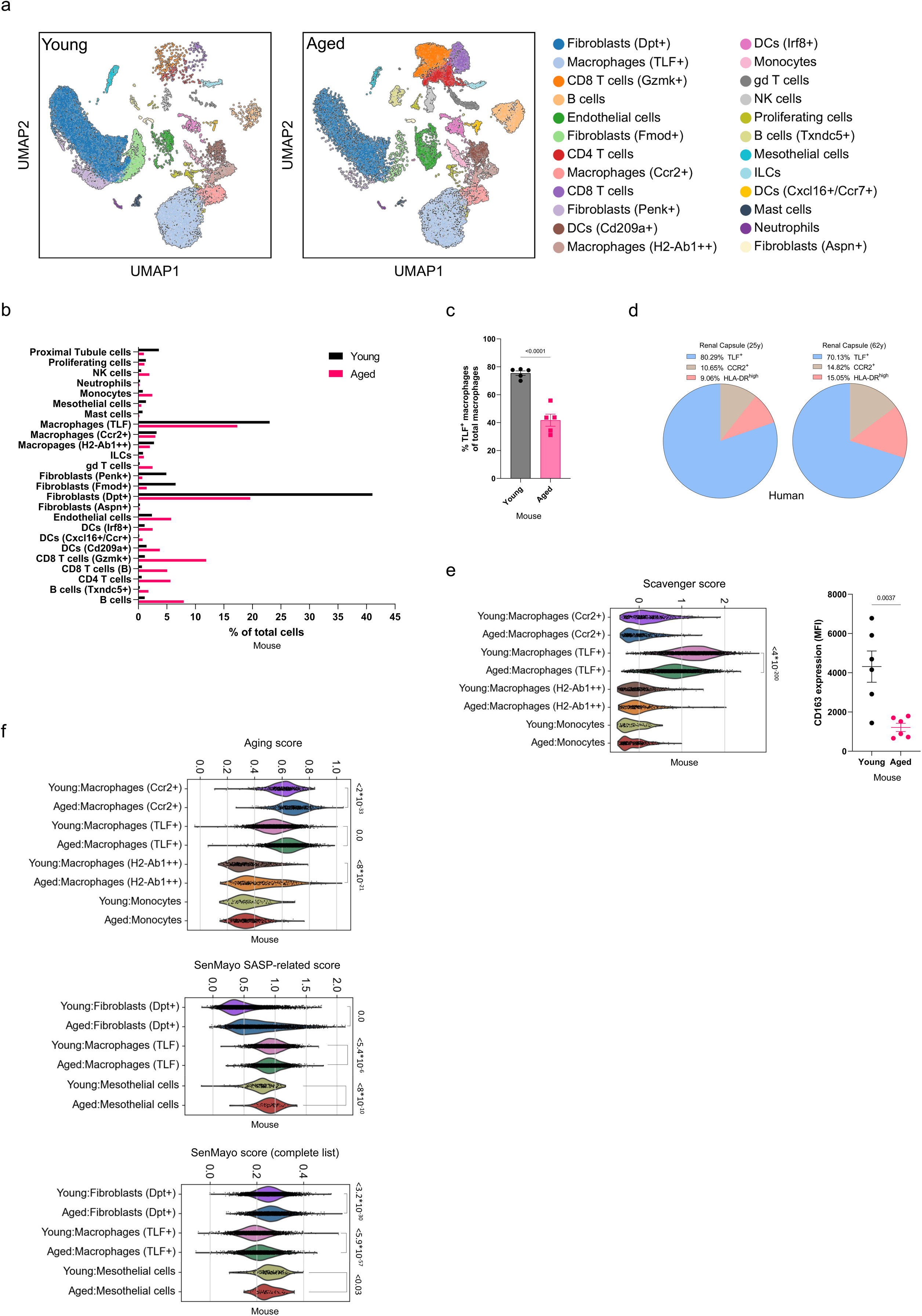
scRNA-Seq identifies complex and cell type specific age-related changes in the renal capsule. (**a**) scRNA-Seq data UMAP of young (3 months) and aged (24 months) mouse renal capsule cells (for each UMAP, n=2 samples, each sample was prepared by pooling renal capsules of 4 mice). (**b**) Percentage of each cell type in young and aged renal capsules (mouse) taken from the UMAP counts in Figure 5a. (**c**) Flow cytometry analysis of TLF^+^ macrophage percentage in young and aged mice (n=6,6, Unpaired t-test). (**d**) Pie chart of macrophage subset proportions in human donors: 25-year-old (as shown in Figure 1) and 62-year-old human donors. (**e**) Scavenger score (**Supplementary Table 3**) of renal capsule myeloid cells (scRNA-Seq data; Mann-Whitney U test), and CD163 expression in young and aged renal capsule TLF^+^ macrophages (n=6,6, unpaired t-test). (**f**) ‘Aging score’ and SenMayo score (SASP-related and complete list) violin plots (**Supplementary Table 3**) of selected cell types, scRNA-Seq data (Mann-Whitney U test).

Since macrophages were the predominant immune population in the renal capsule, we next focused on whether there were age-related changes in macrophage subset composition in the murine renal capsule. Our scRNA-Seq analysis showed a reduction in the number of renal capsule TLF^+^ macrophages in aged mice (**Figure 4b**) and this was confirmed by flow cytometry (**Figure 4c**, **Supplementary Figure 9d**). This reduction was even greater when the increased weight or volume of the aged mouse kidney was taken into account (**Supplementary Figure 9e-g**). With the diminished numbers of TLF^+^ macrophages with age, the average percentages of CCR2^+^ and MHC-II^high^ macrophage subsets were increased (**Supplementary Figure 9h**). Analysis of murine RCAM bulk RNA-Seq further validated this loss of the TLF^+^ subset (**Supplementary Figure 9i**).

We confirmed a similar shift in humans. A 62-year-old donor had a lower percentage of renal capsule TLF^+^ macrophages when compared to a 25-year-old donor (**Figure 4d**). Using samples from multiple donors, we identified that the average percentage of MHC-II^high^ macrophages in the renal capsules of older individuals was higher (**Supplementary Figure 9j**). Next, to elaborate the cell biology of monocyte and macrophages in the aged renal capsule, we investigated function-associated changes by using the metric devised by Sanin and collegues^15^. The aged TLF^+^ macrophages had little differences in ‘phagocytic score’ and ‘inflammatory score’ (**Supplementary Figure 10a,b**, **Supplementary Table 3**). However, a lower scavenger score and a reduction in the expression of the scavenger receptor CD163 was apparent in aged renal capsule TLF^+^ macrophages (**Figure 4e, Supplementary Figure 10c,d**). This suggests that aging may result in the reduction of the scavenging capacity of renal capsule TLF^+^ macrophages.

Cell senescence is an important component of aging, and many studies show that senescent cells and the senescence-associated secretory phenotype (SASP) are associated with detrimental outcomes in the kidney and other organs^22–24^. SASP proteins, secreted by senescent cells, can influence the tissue cell composition, for example, by attracting pro-inflammatory cells^25^. We sought to determine whether cells in the aged renal capsule might have a senescent phenotype. Using an ‘aging score’ based on known aged-related markers, and a previously described murine gene set signature score that predicts senescence-associated pathways, SenMayo^26^, we queried our scRNA-Seq data (**Figure 4f**, **Supplementary Figure 11a**, **Supplementary Table 3**). The ‘aging score’ was highest in macrophages (*Ccr2^+^*), and TLF^+^ macrophages. In TLF^+^ macrophages this result was mediated by the increased expression of *Ccl8*, *Cd209b*, *Cd209d*, *Cd209f*, *Cd209g*, *Cd63*, *Cxcl13*, *Trem2*, and *Vcam1* (**Supplementary Figure 11b**), genes that had a similar shift of expression in aged RCAMs (**Supplementary Figure 11c**). *Ccl8* expression was also higher in aged CCR2*^+^* and MHC-II^high^ macrophages (**Supplementary Table 5**), further suggesting its importance in inflammation of the aged tissue. When using the significantly upregulated SASP-related gene score (murine, 20 gene list), aged *Dpt^+^*fibroblasts, and mesothelial cells scored highly, and aged TLF^+^ macrophages scored lower (**Figure 4f**, **Supplementary Figure 12a**). When the complete SenMayo gene list was used (**Supplementary Figure 12b**, **Supplementary Table 3**), aged TLF^+^ macrophages had a higher score, mostly affected by the increased expression of *Axl*, *Ccl8*, *Cstb*, *Mif* (macrophages migration inhibitory factor), and *Plau* (plasminogen activator, urokinase) (**Supplementary Figure 12c**). Notably, *Cxcl12* expression was highest in aged fibroblasts (*Dpt^+^*) (**Supplementary Figure 12d**). Both scores suggested a higher senescence-associated phenotype in fibroblasts (*Dpt^+^*) with conflicting results in TLF^+^ macrophages.

Using a ‘SASP score’ (**Supplementary Table 3**, based on^27^), two subsets of renal capsule CD8 T cells had the highest score (**Supplementary Figure 13a**). *Ccl5* was the major contributor to this result (**Supplementary Figure 13b**). Additionally, *Mif* chemokine/cytokine was highly expressed in lymphocytes (**Supplementary Figure 13c**), especially in aged CD8 T cells (*Gzmk^+^*), suggesting a role in stimulating innate immune responses and/or recruitment of leukocytes to the capsule. Together, these results suggested that CD8 T cells, *Dpt^+^* fibroblasts, and TLF^+^ macrophages change during renal capsule aging.

Ligand–receptor pairs have recently been used to infer intercellular communication from the expression of their genes^28^. Using the cell-cell communication analysis CellChat developed by Jin and colleagues^29^, we explored signaling ligand-receptor interactions between the cells in the renal capsule. This revealed an overall increased numbers of interactions and higher interaction strengths between the three cell types in the aged renal capsule (**Supplementary Figure 14a**; **Supplementary Table 6**), with increased outgoing and incoming signaling patterns (**Supplementary Figure 14b**). These results support our previous data since they suggest the increased expression of certain ligand-receptor marker combinations in the aged group. For instance, aged TLF^+^ macrophages showed more CXCL-related signaling with CD8 T cells (**Supplementary Figure 14c**; **Supplementary Table 6**). Only the aged cell networks had IL-1 signaling pathways (**Supplementary Figure 14d**), a cytokine with pro-inflammatory properties. Collectively, we propose that aging negatively affects macrophages, fibroblasts, and CD8 T cells in the renal capsule, and together, these cells could promote inflammation, fibrosis, and cell senescence (**Supplementary Figure 15**). Further studies are needed to establish the interactions between these and other cells in the aged renal capsule.

## Discussion

In this study, we characterized cells of the renal capsule following various perturbations and conditions. We found that the renal capsule encompasses a variety of cell types, mostly macrophages and stromal cells. Importantly, compared to the kidney parenchyma, we identified that the mouse and human renal capsule is a unique TLF^+^ macrophage niche. TLF^+^ macrophages are mostly maintained via self-renewal with little monocyte input, embryonically derived, and are the most conserved subset across tissues^14^. Compared to the kidney’s macrophage subset composition, TLF^+^ macrophages have a much higher proportion in organs such as the brain, liver, lung and heart^14^. In murine fetal development, however, TLF^+^ macrophages are much more abundant in the kidney parenchyma^14, 30^. This explicit transition in the kidney may point to the importance of TLF^+^ macrophages in renal development and maturity, highlighting the potential role of these cells when homeostasis is compromised in adulthood. By comparing the properties of RCAMs with their kidney counterparts, we showed that these cells have distinctive transcriptional and cell surface phenotypes, further supporting their unique properties. The distinct anti-inflammatory phenotype and ECM-related transcriptional signature of RCAMs suggests they have a role in tissue remodeling and healing. In the aorta, for example, macrophages (Lyve-1^+^) were shown to maintain arterial tone via hyaluronic-mediated regulation of ECM deposition, possibly affecting its stiffness^31^. RCAMs and TLF^+^ macrophages of the renal capsule may share many properties and cell markers with other serosal macrophages such as peritoneal macrophages, heart or liver serosa macrophages, but may have their own specialized traits that support their function in the kidney. Additional experimental data may reveal such properties and functions in various perturbations.

Focusing on the immune cell dynamics of the renal capsule in male and female mice, during gestation, in obesity, and in renal injury (NTN), we identified that the renal capsule is dynamic, and can respond to changes in the kidney parenchyma. For example, the increased proportion of RCAMs during gestation (E17) may be indicative of their role in renal changes during pregnancy^16^. In the NTN model, an inflammatory surge is typically observed in the first 1-7 days post injection of nephrotoxic serum^32^, which is accompanied by organ expansion. Interesting changes in macrophages were noted as late as 100 days post injection. Nevertheless, the dynamics and composition of immune cells in the renal capsule changed with glomerular injury, suggesting a crosstalk between the different parts of the organ.

Within the aged renal capsule, we identified a heightened inflammatory and fibrotic state, evident by increased immune cell counts. Aging changed the transcriptional phenotype of RCAMs to a more pro-inflammatory pro-fibrotic state, with higher expression levels of cytokines, chemokines, and ECM-related genes. Aged mouse and human renal capsules had less TLF^+^ macrophages and more MHC-II^high^ and CCR2^+^ macrophages. While the MHC-II^high^ subset receives modest monocyte contribution and is not continually replaced, the CCR2^+^ subset is almost entirely replaced by monocytes and is therefore more dynamic. Interestingly, chronic kidney disease is associated with increased numbers of HLA-DR^high^ circulating pro-inflammatory intermediate monocytes, increased migration toward CCL5, and adhesion to primary endothelial cell layers^33^. In the aged mouse brain, it has been reported that parenchymal border macrophages comprise of less Lyve-1^+^ cells and more MHC-II^+^ cells^11, 34^. In the heart, CCR2^+^ macrophage density is correlated with adverse outcomes^35^.

Aged renal capsule TLF^+^ macrophages had a higher ‘aging score’, expressed more age-related and inflammatory markers, suggesting that they are negatively affected by aging and contribute to tissue inflammation. Our scRNA-Seq analysis of young and aged renal capsules suggested that aged TLF^+^ macrophages have reduced scavenger capacity. A reduced scavenger capacity was also reported in aged parenchymal border macrophages^11^, and depletion of TLF^+^ macrophages is associated with lower cellular resilience and regenerative capacity in the heart^36, 37^ and skin^38^, suggesting that these cells are important for organ homeostasis and regeneration.

Macrophages and fibroblasts have important interactions in tissues^39–41^, and age has been shown to negatively affect fibroblast phenotype and function^42, 43^. We explored age-related changes in cell interactions in the renal capsule, focusing on TLF^+^ macrophages, fibroblasts and CD8 T cells, as those populations showed most changes in cell counts in the aged renal capsule. Senescence is known to drive the age-related phenotype, but the definition and characteristics of senescence likely varies between different tissues and cell types. We explored using various scoring systems to score cells with age-related or senescence-associated phenotypes. Using the SenMayo^26^ score (top genes and complete list), aged renal capsule fibroblasts (*Dpt^+^*) had the highest scores. In the aged renal capsule, despite the higher SenMayo scores, fibroblasts did not score highly in the ‘SASP score’, suggesting that age-related changes in kidney parenchymal fibroblasts^27^ are different from fibroblasts in the renal capsule. *Gzmk*^+^ CD8^+^ T cells have been shown to accumulate in aged tissues, including the kidney^27^. We detected increased *Gzmk*^+^ CD8^+^ T cells in the renal capsule and scRNA-Seq analysis showed that they have a higher ‘SASP score’, likely due to IL-1 signaling, which we identified in the aged cell networks using cell-cell communication pathway analysis (CellChat^29^). Additionally, the increased expression of *Ccl8* and *Cxcl12* by macrophages and fibroblasts respectively, likely further contributes to the pro-inflammatory state of the aged renal capsule. Other cell interaction and signaling analysis methods (e.g., NicheNet^44^) could reveal additional insight on this environment in aging. Collectively, we provide an in-depth analysis of the age-related changes that accompany the aged renal capsule. We speculate that the interplay between key players, CD8 T cells, stromal cells, and macrophages contributes to cell subset shifts, increased age-related inflammation, fibrosis, and poor tissue outcome. These data may further our understanding of age and disease-associated changes in the renal capsule, and could provide valuable insight for future studies of serosal tissues in aging, cell senescence, and other conditions.

## Methods

### Animals

C57BL/6J male mice (Jax: 000664), 3-4 months of age (Young) or 22-24 months of age (Aged), weighing at least 20 g or 30-35 g respectively, were used. C57BL/6J female mice (Jax: 000664) were impregnated and used at E17 or 28 days post-delivery. B6.V-Lep^(ob)/J^ Hom Lep (ob) FAT (Jax: 000632) and controls were used at 12 months of age to present a significant obese phenotype (confirmed by weight, which was above 45-50 grams in all mice of the obese group). All animals were housed at Genentech under specific pathogen-free conditions with free access to chow and water and 12/12 hr light/dark cycle. Sample sizes were chosen based on similar experiments that have been previously published. All animal procedures were conducted under protocols approved by the Institutional Animal Care and Use Committee at Genentech, and were performed in accordance with the Guide for the Care and Use of Laboratory Animals.

### Mouse renal capsule dissociation to single cell suspensions

Kidneys were extracted from the euthanized animal without adjacent muscle or adipose tissue. Renal capsules were gently stripped from the kidney with fine forceps, without adipose tissue or renal parenchyma. It is possible that our preparations of the capsule contain some cells associated with the surface of the parenchyma, it is unlikely that deeper cells are present. Tissues were transferred to 1 ml digestion buffer with Liberase^TM^ (1.5 U/ml; 5401127001, Millipore Sigma) and DNaseI (50 U/ml; D4527, Millipore Sigma) in RPMI-1640. Tissues were shredded using scissors and incubated for enzymatic digestion (37⁰C, 800 rpm, 15 min); samples were then sieved via a 100 μm then 70 μm MACS^®^ SmartStrainers (130-098-463, 130-098-462, Miltenyi Biotec) and 10 ml ice-cold 10% fetal bovine serum (FBS) in RPMI-1640 were added to stop digestion. Samples were kept on ice for further processing.

### Mouse kidney dissociation to single cell suspensions

Following extraction of renal capsules from kidneys, kidneys were minced using a razor blade, then transferred to 5 ml digestion buffer with Liberase^TM^ (1.5 U/ml; 5401127001, Millipore Sigma) and DNaseI (100 U/ml; D4527, Millipore Sigma) in RPMI-1640, and incubated for enzymatic digestion (37⁰C, 800 rpm, 20 min). Following digestion, samples were sieved via a 100 μm and a 70 μm MACS^®^ SmartStrainers (130-098-463, 130-098-462, Miltenyi Biotec), and 30 ml ice-cold 10% FBS in RPMI-1640 were added to stop digestion. Samples were kept on ice for further processing.

### Human kidney and renal capsule dissociation to single cell suspensions

Human kidneys we stripped from outer adipose tissue. For renal capsules, five randomly selected ∼1 cm^2^ pieces were taken from each human donor kidney, tissues were transferred to 10 ml digestion buffer with Liberase^TM^ (5 U/ml; 5401127001, Millipore Sigma) and DNaseI (0.1 mg/ml; 07900, Stem Cell Technologies) in RPMI-1640. Tissue was shredded using scissors and incubated for enzymatic digestion (37⁰C, 800 rpm, 30 min), then sieved via a 100 μm and 70 μm MACS^®^ SmartStrainers (130-098-463, 130-098-462, Miltenyi Biotec) and 30 ml ice-cold 10% fetal FBS in RPMI-1640 were added to stop digestion. Processing of kidney sample (∼1 cm^2^ piece) was performed as described above. Samples were kept on ice for further processing.

### Flow cytometry

Single-cell suspensions were washed with cell staining buffer twice (300g, 5 min, 4⁰C) and incubated in 100 μl of Cell Staining Buffer (10 ml; 420201, Biolegend) with TruStain FcX™ (1:100; 101320, Biolegend) and True-Stain Monocyte Blocker™ (1:20; 426103, Biolegend) for FC blocking (15 min, 4⁰C); then stained with a mixture of fluorescently labelled anti-mouse antibodies (30 min, 4⁰C; **Supplementary Table 1**) in Cell Staining Buffer. Live-dead cell discrimination was performed using Zombie Aqua™ Fixable Viability Kit (423102, Biolegend) according to manufacturer instructions. Following staining, cells were washed three times (300g, 5 min, 4⁰C) in cell staining buffer and kept in FluoroFix™ Buffer (422101, Biolegend) prior to run in a BD FACSymphony™ A5 Cell Analyzer (BD Biosciences). Analysis was performed using Cytobank (Cytobank.org) or FlowJo™ v10.8 Software (BD Life Sciences).

### Mass cytometry

Due to limited cell numbers from each renal capsule, single-cell renal capsule samples were pooled from several mice (as indicated in each figure legend) for subsequent mass cytometry analysis as follows. Single cell suspensions of the renal capsule were washed with Maxpar Cell Staining Buffer (201068, Fluidigm) twice (300g, 5 min) and incubated in 100 μl of Maxpar Cell Staining Buffer with TruStain FcX™ (anti-mouse CD16/32) Antibody (1:100; 101320, Biolegend) for FC blocking (10 min, 4⁰C). For mass cytometry analysis of renal capsule and kidney cells, samples were barcoded with differently labelled CD45 antibodies and run in the same sample (n=2 pooled mice). After blocking and optional sample barcoding, samples were stained with a mixture of anti-mouse metal-tagged antibodies for surface markers (1:100, 45 min, 4⁰C; **Supplementary Table 2**). Rhodium (1:1000; 201103A, Fluidigm) was added to the cells in the last 15 min of staining. Intracellular staining was performed flowing fixation with 1X Maxpar Fix I Buffer (15 min, room temperature) and two washes with Maxpar Perm-S Buffer (500 g, 5 min) with a mixture of anti-mouse metal-tagged antibodies for intracellular markers (1:100, 30 min, room temperature; **Supplementary Table 2**). Cells were then washed twice with Maxpar Cell Staining Buffer, and fixed in freshly made 1.6% paraformaldehyde (PFA; 50-980-487, Electron Microscopy Sciences) in Maxpar PBS (1 hr, room temperature; 201058, Fluidigm). Prior to sample run on a Helios mass cytometer (Fluidigm), cells were incubated with Cell-ID™ Intercalator-Ir (1:1000, 20 min, room temperature; 201192A, Fluidigm) in Maxpar Fix and Perm Buffer (201067, Fluidigm), washed once in Maxpar Cell Staining Buffer (500 g, 5 min) and then washed twice with Maxpar Water (500 g, 5 min; 201069, Fluidigm). Cell concentration was adjusted to <10^6^ cells/ml in Maxpar Water and EQ Four Element Calibration Beads (201078, Fluidigm) were added for sample normalization. Data were acquired using a Helios Mass Cytometer (Fluidigm) at approximately 500 events/sec. Data were normalized in CyTOF Software (Fluidigm) and uploaded to Cytobank (Cytobank.org) for analysis.

### Whole mount renal capsule staining and imaging

Immediately after being euthanized, mice were transcardially perfused with 20 ml ice-cold phosphate buffered saline without calcium and magnesium (PBS; 21-040-CV, Corning) followed by 20 ml ice-cold 4% PFA (15710, Electron Microscopy Solutions) in PBS. Kidneys were extracted, any excess tissue was gently removed, and placed in 4% PFA in PBS for 2 hr at room temperature. Renal capsules were gently stripped from kidneys and washed in PBS three times (15 min, room temperature). Tissues were permeabilized and blocked in blocking buffer with 0.05% Tween 20 (H5152, Promega) and 5% BSA (A3059, Millipore Sigma) in PBS for 1.5 hr at room temperature. Tissues were then stained with the following fluorophore-conjugated anti-mouse antibodies for (1:100, 1.5 hr, room temperature, gentle agitation) in blocking buffer: Iba-1 (78060S, Cell Signaling Technology), F4/80 (123122, Biolegend), CD206 (141710, Biolegend), CD45 (103172, Biolegend), and Lyve-1 (FAB2125G, R&D). At the last 15 min of staining, Hoechst (ab228551, Abcam) was added (1:2000) for nuclei staining. For incubation with unconjugated antibodies (1:200, 1.5 hr, room temperature, gentle agitation), Decorin (ab175404, Abcam), PDGFRa (ab203491, Abcam), tissues were washed in PBS three times (15 min, room temperature) and incubated (45 min, room temperature, gentle agitation) with a secondary fluorescently labelled antibody (A-31573, Invitrogen, 1:500) in blocking buffer. Lastly, tissues were washed in PBS three times (15 min, room temperature), mounted with ProLong™ Diamond Antifade Mountant (P36965, Invitrogen) and sealed with VWR micro cover glass (48404-455, VWR). In slides with DAPI, ProLong™ Gold Antifade reagent with DAPI was added when slide was sealed. Images were taken with Thunder Imaging System (Leica Microsystems).

### Whole kidney tissue staining, clearing and imaging

Immediately after being euthanized, mice underwent transcardial perfused with 20 ml ice-cold PBS and then with 20 ml ice-cold 4% paraformaldehyde (PFA) in PBS. Kidneys were extracted, any excess tissue was gently removed, and placed in 4% PFA in PBS for 24 hr at 4⁰C. Tissue clearing was performed using Ce3D™ Tissue Clearing kit (427702, Biolegend) according to manufacturer instructions with modifications. Briefly, following fixation, kidneys were washed three times for 1 hr at room temperature in Ce3D™ Wash Buffer (427710, Biolegend), then cut in half and transferred to Ce3D™ Permeabilization/Blocking Buffer (48 hr, room temperature, gentle shaking; 427706, Biolegend). The Ce3D™ Permeabilization/Blocking Buffer was discarded and replaced with Alexa Fluor 647 conjugated anti-mouse Iba-1 (1:100, 78060S, Cell Signaling Technology) in Ce3D™ Antibody Diluent Buffer (48 hr, room temperature, with gentle shaking; 427708, Biolegend). Following staining, samples were washed three times in Ce3D™ Wash Buffer (8 hr, room temperature, gentle shaking). Before imaging, Ce3D™ Wash Buffer was removed completely and samples were placed in Ce3D™ Tissue Clearing Solution (12 hr, room temperature, with gentle shaking; 427704, Biolegend). Images were taken with Leica SP8 Confocal System (Leica Microsystems).

### Fluorescence activated cell sorting (FACS)

Mouse renal capsule or kidney single cell suspension samples were washed with cell staining buffer twice (300g, 5 min, 4⁰C) and incubated in 100 μl of Cell Staining Buffer with TruStain FcX™ (anti-mouse CD16/32) Antibody (1:100; 101320, Biolegend) and True-Stain Monocyte Blocker™ (1:20; 426103, Biolegend) for FC blocking (15 min, 4⁰C); then stained with a mixture of fluorescently labelled anti-mouse antibodies in Cell Staining Buffer (30 min, 4⁰C): BV785 conjugated CD45, PerCP-Cy5.5 conjugated CD11b, BUV395 conjugated F4/80, PE conjugated CD4, PE conjugated CD19, PE conjugated CD8, PE conjugated Gr-1. Live-dead cell discrimination was performed using Zombie Aqua™ Fixable Viability Kit (1:1000, 423102, Biolegend) according to manufacturer instructions. Macrophages were gated and sorted according to singlets/Zombie Aqua^neg^/CD45^+^/PE^neg^/CD11b^+^/F480^+^. Samples were sorted using BD FACSAria™ Fusion Cell Sorter (BD Biosciences).

### Nephrotoxic nephritis (NTN) mouse model

NTN mouse model was performed by an intravenous tail vein injection of 5 μl/g mouse of sheep anti-rat glomerular serum (nephrotoxic serum (NS); Probetex), as previously described^32^. Briefly, injection of NS causes an acute inflammatory response and proteinuria (verified by Uristix, reagent strips for urinalysis; Siemens) that peak after 24 hours post-injection and decrease over time. Thirty-five days later, glomerulosclerosis and fibrosis are evident, indicating a sustained renal injury. Proteinuria and kidney inflammation occur over the first 1-7 days and are largely resolved by day 35. At this time point, there is no change in kidney function (normal glomerular filtration rate (GFR)). This treatment was performed on C57BL/6J male mice to minimize experimental variability.

### Tissue-residing cell analysis

Mice were anaesthetized using a 4% isoflurane and medical air and maintained in 1–2% isoflurane during the following procedures. At the end of each procedure, mice were euthanized by cervical dislocation or CO_2_ inhalation.

Perfusion-based analysis: mice were transcardially perfused with 20 ml ice-cold PBS (21-040-CV, Corning). Kidneys were extracted, any excess tissue was gently removed, and renal capsules were stripped from kidneys. Renal capsules were processed into single cell suspensions as mentioned previously, stained and analyzed using flow cytometry.

Intravenous (i.v.) injection-based analysis: i.v. injection of an anti-CD45 antibody was performed as previously described^45, 46^ with some modifications. Briefly, mice were subjected to i.v. injection of APC conjugated anti-mouse CD45 (cat# 103112; clone 30-F11; Biolegend) at 10 µg per mouse; 5 minutes later, mice were immediately euthanized, kidneys were extracted and renal capsules were stripped for further processing and analysis by flow cytometry. Blood was taken to validate labelling by the injected antibody and negative control mice were injected with PBS.

### Bulk RNA sequencing (bulk RNA-Seq) and analysis

For bulk RNA sequencing, cells were sorted directly into RNeasy Lysis Buffer (Qiagen). RNA was extracted and isolated from samples using RNeasy Micro Kit (74004, Qiagen) according to manufacturer instructions. **Regular input RNA-Seq**: total RNA was quantified with Qubit RNA HS Assay Kit (Thermo Fisher Scientific) and quality was assessed using RNA ScreenTape on 4200 TapeStation (Agilent Technologies). For sequencing library generation, the Truseq Stranded mRNA kit (Illumina) was used with an input of 100 nanograms of total RNA. Libraries were quantified with Qubit dsDNA HS Assay Kit (Thermo Fisher Scientific) and the average library size was determined using D1000 ScreenTape on 4200 TapeStation (Agilent Technologies). Libraries were pooled and sequenced on NovaSeq 6000 (Illumina) to generate 30 million single-end 50-base pair reads for each sample. **Low input RNA-Seq**: total RNA was quantified with Qubit RNA HS Assay Kit (Thermo Fisher Scientific catalog#: Q32852) and quality was assessed using RNA ScreenTape on 4200 TapeStation (Agilent Technologies catalog#: 5067-5576). cDNA library was generated from 2 nanograms of total RNA using Smart-Seq V4 Ultra Low Input RNA Kit (Takara catalog#: 634894). 150 picograms of cDNA was used to make sequencing libraries by Nextera XT DNA Sample Preparation Kit (Illumina catalog#: FC-131-1024). Libraries were quantified with Qubit dsDNA HS Assay Kit (Thermo Fisher Scientific catalog#: Q32851) and the average library size was determined using D1000 ScreenTape on 4200 TapeStation (Agilent Technologies catalog#: 5067-5582). Libraries were pooled and sequenced on NovaSeq 6000 (Illumina) to generate 30 million single-end 50-base pair reads for each sample. Data were analyzed and visualized using *Partek^®^ Flow^®^* (https://www.partek.com/partek-flow/).

### Single cell RNA sequencing (scRNA-Seq) and analysis

Processed samples were immediately sorted (FACS) according to size (FSC-A) and complexity (SSC-A), to minimize cell debris and achieve higher sample purity and viability (>85%). Sample processing for scRNA-seq was done using Chromium Next GEM Single Cell 3’ Kit v3.1 (1000269, 10X Genomics) according to manufacturer’s instructions. Cell concentration was used to calculate the volume of single-cell suspension needed in the reverse transcription master mix, aiming to achieve approximately 8000 cells per sample. cDNA and libraries were prepared according to manufacturer’s instructions (10X Genomics). Sequencing reads were processed through 10X Genomics Cell Ranger (v.3.1.0) and aligned to GRCm38. We used Scanpy^47^ for downstream data analysis. Cells with fewer than 400 genes expressed or more than 5,000 genes expressed were filtered out. Genes that were expressed in less than 50 cells where removed. Cells with 10% or more mitochondrial content were also removed. Scrublet^48^ was used to remove doublets. Harmony^49^ was used to harmonize the data from the different samples. Clustering analysis was performed using Leiden clustering with a resolution set to 0.8. Initial cluster annotation was performed using SingleR^50^. Inference of cell-cell communication was performed using CellChat^29^. Gene set scores were calculated using scanpy’s score_genes function.

### Histology and pathology analysis of mouse and human samples

Tissues used for histologic analysis and immunohistochemistry (IHC) were routinely fixed in formalin, paraffin embedded, and sectioned at approximately 4-5 µm. Mouse tissues were histologically scored in a random, blinded manner on hematoxylin and eosin-stained sections. The scoring system included assessments of inflammation (0: no significant inflammatory infiltrates; 1: 1-2 small leukocyte aggregates (less than approximately 10 cells or scattered leukocytes associated with adjacent tubular atrophy); 2: 3-5 small leukocyte aggregates or 1 larger aggregate; 3: 6-10 small leukocyte aggregates or 2-3 larger aggregates; 4: >10 small leukocyte aggregates or >3 larger aggregates) and subcapsular tubular atrophy and fibrosis (0: no subcapsular tubular atrophy; 1: 1-2 foci of subcapsular tubular atrophy; 2: 3-5 foci of subcapsular tubular atrophy; 3: 6-10 foci of subcapsular tubular atrophy; 4: >10 foci of subcapsular tubular atrophy). These scores were summed to generate a final score.

IHC was performed for collagen 3 (Southern Biotechnology goat polyclonal, 7.5 µg/ml), Pax5 (Abcam rabbit monoclonal antibody EPR3730(2), 0.134 µg/ml), CD3 (Thermo Scientific rabbit monoclonal antibody SP7, 0.134 µg/ml), IBA1 (Abcam rabbit monoclonal antibody EPR16588, 0.25 µg/ml), CD68 (Dako mouse monoclonal antibody KP1, 0.1 µg/ml), fibroblast activation protein (FAP, Abcam rabbit monoclonal antibody EPR20021, 2.5 µg/ml), and vimentin (Cell Signaling rabbit monoclonal antibody D21H3, 0.45 µg/ml). Collagen 3, Pax5, CD3, CD68, and FAP IHC was performed on the Ventana Discovery XT platform with CC1 standard antigen retrieval, and OmniMap detection with a diaminobenzidine chromogen. Bovine serum albumin was used to block immunoglobulin binding for the Pax5, CD3, and FAP assays. TNB was used for the CD68 assay. IBA1 was performed similarly on the Discovery XT platform with OmniMap detection, and bovine serum albumin block, except CC1 mild was used for antigen retrieval. Vimentin immunolabeling was performed on a Dako autostainer with Target retrieval and hydrogen peroxidase, ScyTek block, and donkey serum for blocking non-specific labeling. Immunolabeling was detected with ABC-Peroxidase Elite (Vector Labs) and diaminobenzidine. Positive controls included multiple mouse tissues for collagen 3, spleen for Pax5 and CD3, human brain for IBA1, human tonsil for CD68, human urothelial carcinoma for FAP, and normal human liver and kidney for vimentin. Isotype-matched were also included in each run.

Collagen 3 labeling was scored in a random and blinded manner according to the following matrix: 0, no increased fibrosis; 1, 1-2 fibrotic foci; 2, 2-3 fibrotic foci; 3, >5 fibrotic foci or confluent fibrosis; 4, extensive capsular fibrosis and remodeling. The number of Pax5 and CD3 positive cells in the capsule were counted in 1 sagittal section of each kidney.

### Human Tissue Procurement

All human tissue samples were supplied by Donor Network West. Donor Network West received IRB approval of research, appropriate informed consent of all subjects contributing biological materials, and all other authorizations, consents, or permissions as necessary for the transfer and use of the biological materials for research at Genentech.

Fresh kidney tissue from a total of 7 unique male donors were procured post-mortem. Donors ranged in age between 12 to 66 years and were of Caucasian or Latino descent. Segments of tissue were placed into RPMI/10%FBS and 4% PFA and transferred to Genentech. A portion of each tissue in RPMI/10%FBS was fixed in formalin, processed and embedded. Sections were created and stained with Hematoxylin and Eosin (H&E) for review of morphology.

### Statistical analyses and reproducibility

Data are presented as mean ± s.e.m. Experiments were repeated independently at least two times (biological replicates). Statistical significance was determined using two-tailed unpaired t-test (nonparametric) or Mann-Whitney U test when comparing two independent groups. For comparisons of multiple factors, one-way or two-way analysis of variance (ANOVA) with appropriate multiple-comparisons tests were used. Statistical analyses were performed using Prism 9.5 (GraphPad software). *P values* are provided within figures. Approximate renal capsule surface area (SA) was calculated according to the ellipsoid formula: SA = 4Π[((ab)1.6+(ac)1.6+(bc)1.6)/3](1/1.6) when a,b,c are the measured axis/radius per kidney.

### Data availability

The data that support the findings of this study are available upon reasonable request.

## Acknowledgements

We thank and our Genentech colleagues in the Research Biology, Laboratory Animal Resources, Microscopy and Pathology Departments for their support in this study. We thank Eric Jang and Will Park (Human Tissue Lab) who helped with accessioning of human samples. Genentech hereby expresses its thanks for the cooperation of Donor Network West and all of the organ and tissue donors and their families, for giving the gift of life and the gift of knowledge, through their generous donations. This work was supported by Genentech.

## Contributions

B.K. conceived the project, designed and performed experiments, analyzed and interpreted the data, created the figures and wrote the manuscript. S.A. performed animal experiments, assisted in data interpretation and revised the manuscript. M.C-T assisted in animal experiments. R.M. assisted in scRNA-Seq experiments. T.H. assisted in mass cytometry experiments and cell sorting. H.M, S.D. Y.L. Z.M. supported bulk RNA-Seq and scRNA-Seq. C.C. supported light microscopy imaging. C.V., L.R, C.H. and W.E. - procurement and processing of human samples. C.J. and J.J. - histology monitors. D.D. - mouse IHC. M.D. - human IHC. A.M. supported scRNA-Seq data processing. J.D.W. - pathology analysis and data interpretation, and revised the manuscript. S.D performed scRNA-seq data analysis and revised the manuscript. A.S. oversaw data interpretation, and revised the manuscript.

## Supplementary data legend

**Supplementary Figure 1: Supporting cytometry and imaging data for** Figure 1. (**a**) Gating strategy for live single cells using mass cytometry as described in Figure1a,b. (**b**) Percentage of each cell type as calculated according to the opt-SNE analysis in Figure 1b. (**c**) Opt-SNE plots for various cell markers used to determine cell identity in Figure 1b. (**d**) Histograms of all cell markers used in mass cytometry, indicating marker expression levels for each cell population in Figure 1b. (**e**) Illustration of the experimental procedure for whole mount renal capsule staining and imaging with representative images. Created with BioRender.com. (**f**) Representative immunolabeling (Iba-1, CD68, FAB, Vimentin – right) imaging of human renal capsules. Normal - top, subcapsular inflammation - bottom. These samples represent clinical variability and were used only as an example for healthy and inflamed human renal capsules. (**g**) Illustration of the experimental procedure for tissue clearing. Created with BioRender.com.

**Supplementary Figure 2: Blood cell contamination analyses.** Flow cytometry analyses of: (**a**) renal capsules from perfused and non-perfused (control) mice (2-way ANOVA); (**b**) renal capsule cells taken from PBS (negative control) or anti CD45-APC injected mice, and representative blood samples of anti CD45-APC injected mice (positive control) (n=2,5,2) with representative histograms of each condition or tissue. (**c**) Red blood cell (RBC) staining (anti TER-119) of renal capsule cells (n=4).

**Supplementary Figure 3: Bulk RNA-Seq data of renal capsule associated macrophages (RCAMs) and kidney macrophages**. (**a**) PCA plot according to bulk RNA-Seq data of samples sorted for RCAMs (n=4) or kidney macrophages (n=4) as indicated in Figure 1e. (**b**) Heatmap of top differentiating genes. (**c**) *Gata6* normalized expression in RCAMs (bulk RNA-Seq data (n=4)), peritoneal macrophages and subcutaneous lymph node lymphatic endothelial cells (taken from ImmGene^51^).

**Supplementary Figure 4: Mass cytometry analysis of RCAMs and kidney macrophages**. (**a,b**) Mass and flow cytometry gating strategy for RCAMs and kidney macrophages. (**c**) Representative opt-SNE plots of distinguishing markers between RCAMs and kidney macrophages (n=3 pooled mice for each sample).

**Supplementary Figure 5: Supporting scRNA-Seq data**. (**a**) Cell annotated UMAPs and cell percentage without (left) or with (right) proximal tubule cells. The presented UMAPs were generated using cells from four samples (n=4 pooled mice per sample). (**b**) UMAPs of main identification markers grouped by main cell types. UMAPs include young and aged cells as shown in Figure 4, and proximal tubule cells. (**c**) Mki67 expression and cell cycle analysis UMAPs. (**d**) *Csf1* and *Il34* expression UMAP overlay.

**Supplementary Figure 6: Supporting bulk RNA-Seq and flow cytometry data for RCAMs and TLF^+^ macrophages**. (**a**) Top TLF genes in volcano plot and heatmap of bulk RNA-Seq data generated from sorted RCAMs and kidney macrophages. (**b**) Immunofluorescence imaging of whole mounted renal capsule (Lyve-1, CD45, CD206). (**c**) Histograms depicting the expression of protein markers in mouse (Tim4, Lyve1, FRb, CD44, CD206, CD169, CD163, CD14, CD11c) and human (TIM4, FRb, CD206, CD163, CD44) RCAMs and kidney macrophages, flow cytometry. (**d**) Gating strategy for mouse and human TLF^+^ macrophages.

**Supplementary Figure 7: RCAMs MHC-II expression in various conditions and representative flow cytometry plots**. Relative MHC-II expression was measured to represent activation state of antigen presentation in RCAMs taken from: (**a**) male and female C57BL/6J mice (n=5,5, unpaired t-test), (**b**) pregnant (E17), 28 days post gestation and control female mice (n=6,6,6, two-way ANOVA), (**c**) confirmed by body weight obese (B6.Cg-Lepob/J) and control male mice (1 year-old; n=4,4, two-way ANOVA), (**d**) confirmed by proteinuria (Uristix; left) (**e**) nephrotoxic nephritis (NTN) C57BL/6J male mice (0, 1, 5, 10, 40, and 100 days post nephrotoxic serum (NS) intravenous injection; n=5 per time point, two-way ANOVA). Analyzed by flow cytometry, fold change of control expression (median fluorescence intensity; MFI). (**f**) Representative flow cytometry gating strategy for immune populations in the renal capsule (3- month-old male mouse). (**g-j**) Representative flow cytometry plots for the gestation, obesity, and NTN models. Gate percentage in the dot plot corresponds to the percentage of events within the last indicated parent population (top).

**Supplementary Figure 8: Supporting histology, flow cytometry, and bulk RNA-Seq data of age-related changes in the renal capsule**. (**a**) Histology scores for inflammation, tubular atrophy, fibrosis, and collagen 3, corresponding to the analysis shown in Figure 3a and used to determine the overall histology score. (**b**) T and B cell counts in renal capsule-associated cells of 6, 12, 18, 24-month-old mice, according to CD3 and PAX5 immunohistochemistry. (**c**) Fold change of myeloid and lymphoid cells (2-way ANOVA, adjusted p value<0.0001, n=4,4) in the renal capsules of young and aged mice. (**d**) Percentage of immune cell types in the young and aged renal capsule (n=4,4, 2-way ANOVA). (**e,f**) PCA plots and heatmap comparisons of bulk RNA-Seq analysis of whole tissue and sorted RCAMs of young and aged mice (n=3,3 and n=4,4 respectively). (**g**) Pathway analysis of top differentially expressed genes (upregulated in aged as shown in (e)).

**Supplementary Figure 9: Supporting data of age-related changes in the renal capsule**. (**a**) Overlay UMAP of young and aged renal capsule cells and (**b**) # of cells per cell type as shown in Figure 4a. (**c**) Red blood cell (RBC) staining (anti TER-119) of renal capsule cells from young and aged mice (n=3,3, unpaired t-test; flow cytometry), indicating that the observed increased cell counts in the aged group were independent of blood contamination. (**d**) Number of TLF^+^ macrophages in young and aged renal capsules (n=5,5, unpaired t-test). (**e**) Representative image of young (3 months) and aged (24 months) mouse kidneys. (**f**) Kidney mass (n=5,5, unpaired t-test) and number of TLF^+^ macrophages per kidney mass (n=5,5, unpaired t-test). (**g**) Approximate renal capsule surface area and number of TLF^+^ macrophages per calculated surface area (n=5,5, unpaired t-test). (**h**) Percentages and counts of CCR2^+^ and MHC-II^high^ macrophages of total macrophages (n=5,5, unpaired t-test). (**i**) TLF-associated genes of young and aged RCAMs, bulk RNA-Seq analysis. (**j**) Percentage of HLA-DR^high^ macrophages in young (<37 years, n=3) and aged (>54 years, n=3, unpaired t-test) human renal capsules, analyzed by flow cytometry.

**Supplementary Figure 10: Supporting data of age-related changes in the renal capsule, scRNA-Seq data**. (**a**,**b**) Phagocytosis and inflammatory scores of young and aged myeloid cell types, according to scRNA-Seq data (Mann-Whitney U test). (**c**) Young and aged renal capsule UMAP of scavenger score. (**d**) Bubble plot of scavenger associated genes in young and aged renal capsule TLF^+^ macrophages.

**Supplementary Figure 11: Supporting data of ‘aging score’ related changes in the renal capsule**. (**a**) Young and aged renal capsule overlaid UMAP of ‘aging score’, scRNA-Seq data. (**b**) Bubble plot of ‘aging score’ genes in TLF^+^ macrophages. (**c**) RCAMs bulk RNA-Seq data of selected genes as shown in b.

**Supplementary Figure 12: Supporting data of SenMayo score related changes in the renal capsule.** (**a**) Young and aged renal capsule overlaid UMAP of SenMayo SASP-related score. (**b**) Young and aged renal capsule overlaid UMAP of SenMayo complete list score, with bubble plots of score genes in (**c**) TLF^+^ macrophages and (**d**) fibroblasts (*Dpt^+^*).

**Supplementary Figure 13: Supporting data of ‘SASP score’ related changes in the renal capsule**. (**a**) Young and aged renal capsule overlaid UMAPs of ‘SASP score’, and (**b**) bubble plot of ‘SASP score’ genes in T cells, scRNA-Seq data (Mann-Whitney U test). (**c**) Young and aged renal capsule overlaid UMAPs and violin plot of *Mif* expression.

**Supplementary Figure 14: Supporting data of renal capsule CellChat analysis**. (**a**) Young and aged CellChat plots with interaction weight or strength. (**b**) CellChat heatmap of outgoing and incoming signaling pathways of CD8 T cells, fibroblasts, and macrophages. (**c**,**d**) CellChat signaling pathway network plots used to analyze the scRNA-Seq data of young and aged renal capsule cells. Ligand-receptor details for each pathway are available in^29^).

**Supplementary Figure 15: Schematic description of suggested cell interactions in the renal capsule in aging**.

**Supplementary Table 1**: Detailed information of flow cytometry markers used in this study.

**Supplementary Table 2**: Detailed information of mass cytometry markers used in this study.

**Supplementary Table 3**: Gene lists for scRNA-Seq scores described in this study.

**Supplementary Table 4**: Details regarding histology scores described in this study.

**Supplementary Table 5**: Differential gene analysis of the three macrophage subsets, aged versus young.

**Supplementary Table 6:**CellChat pathway analysis details. Taken from: http://www.cellchat.org/cellchatdb/.

