## Supplementary Figures for "The renal capsule, a vibrant and adaptive cell environment of the kidney in homeostasis and aging"

Supplementary Figure 1: Supporting cytometry and imaging data for Figure 1.

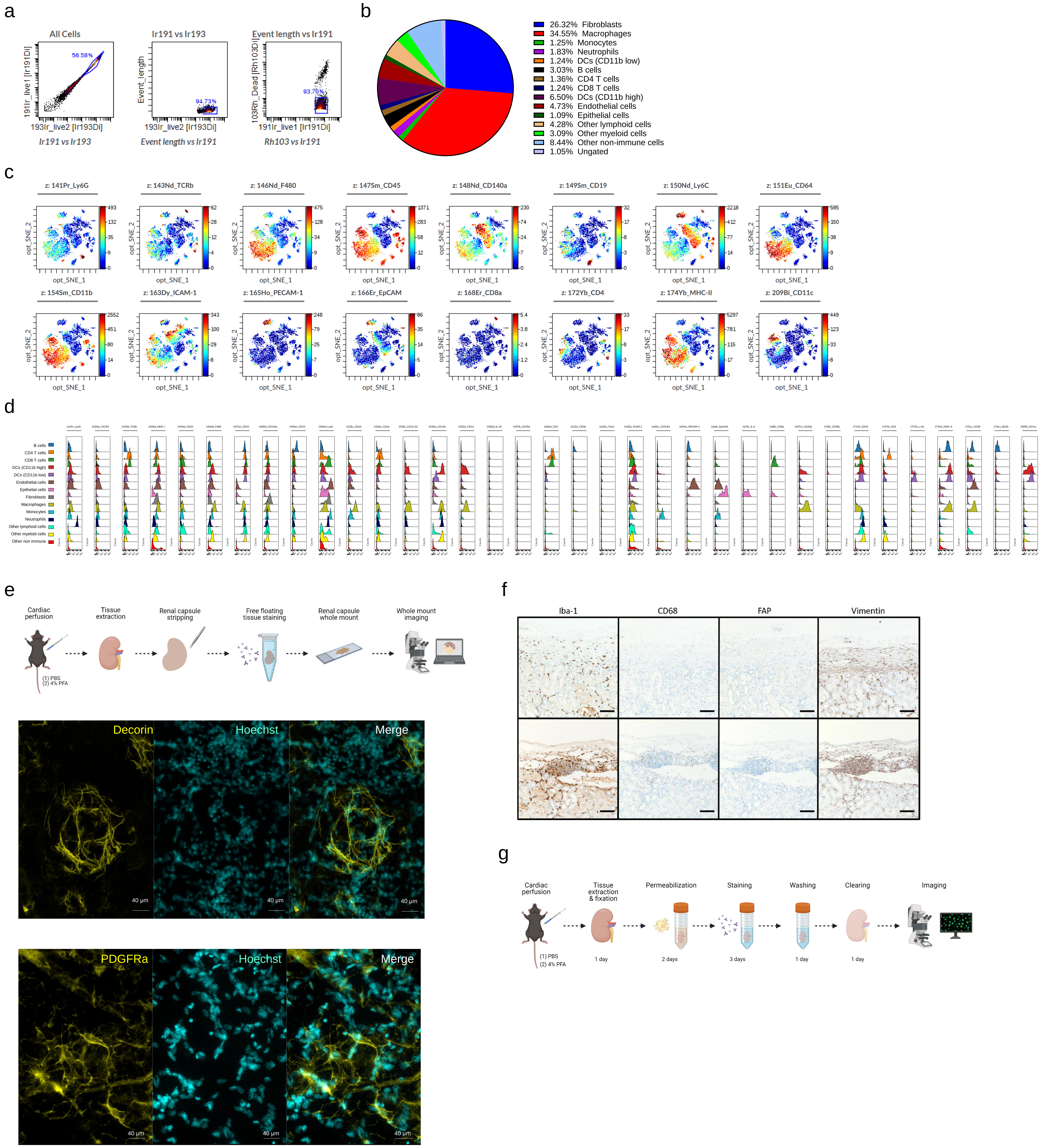

Supplementary Figure 2: Tissue residing cell analysis.

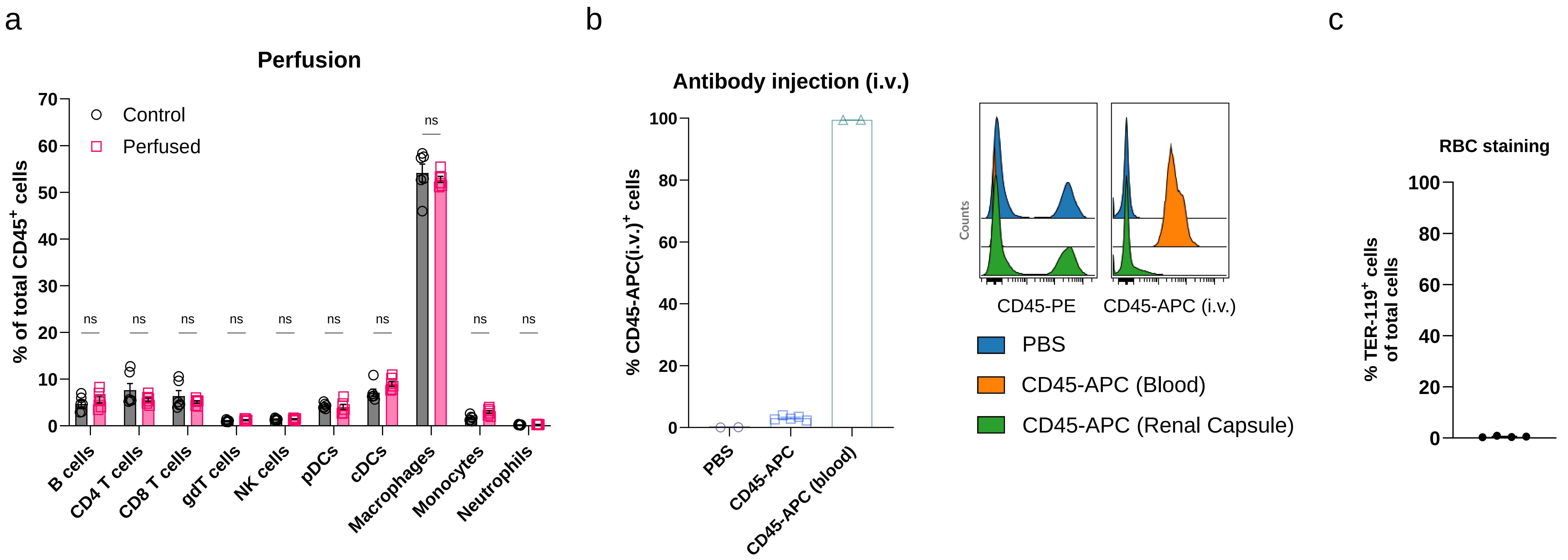

Supplementary Figure 3: Characterization of renal capsule associated macrophages (RCAMs) highlights their distinction from kidney macrophages.

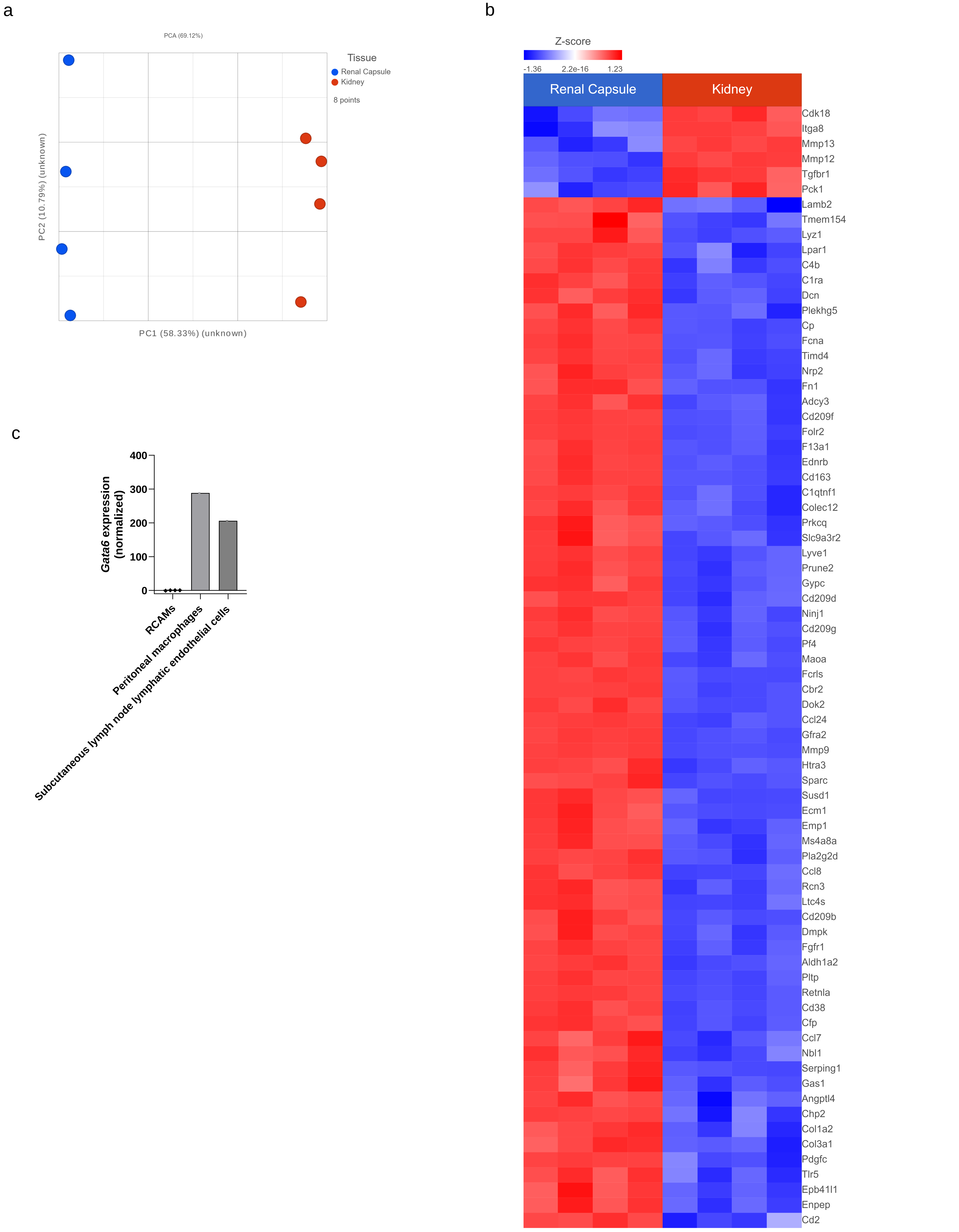

Supplementary Figure 4: Mass cytometry analysis of RCAMs and kidney macrophages.

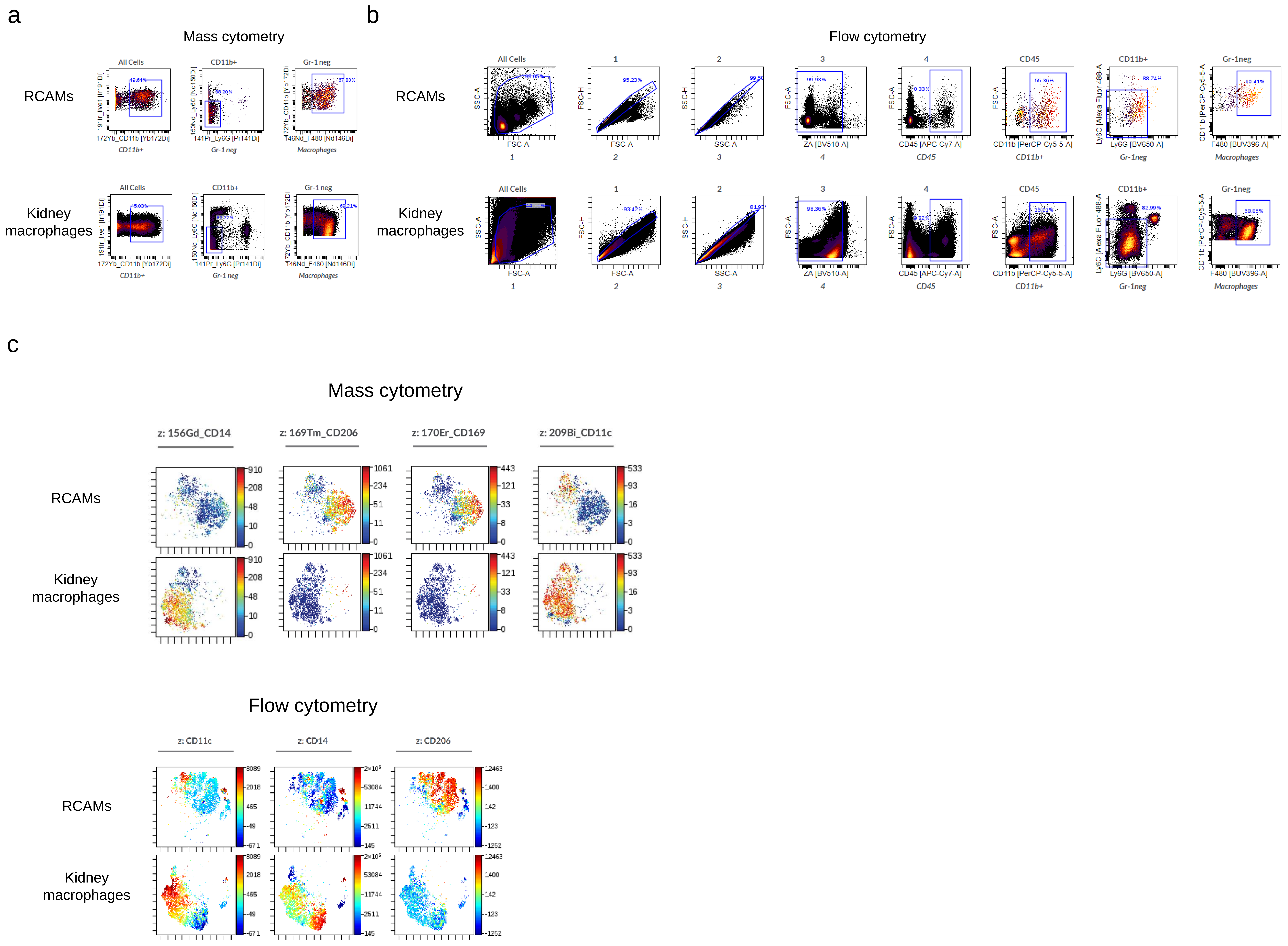

Supplementary Figure 5: Supporting scRNA-Seq data.

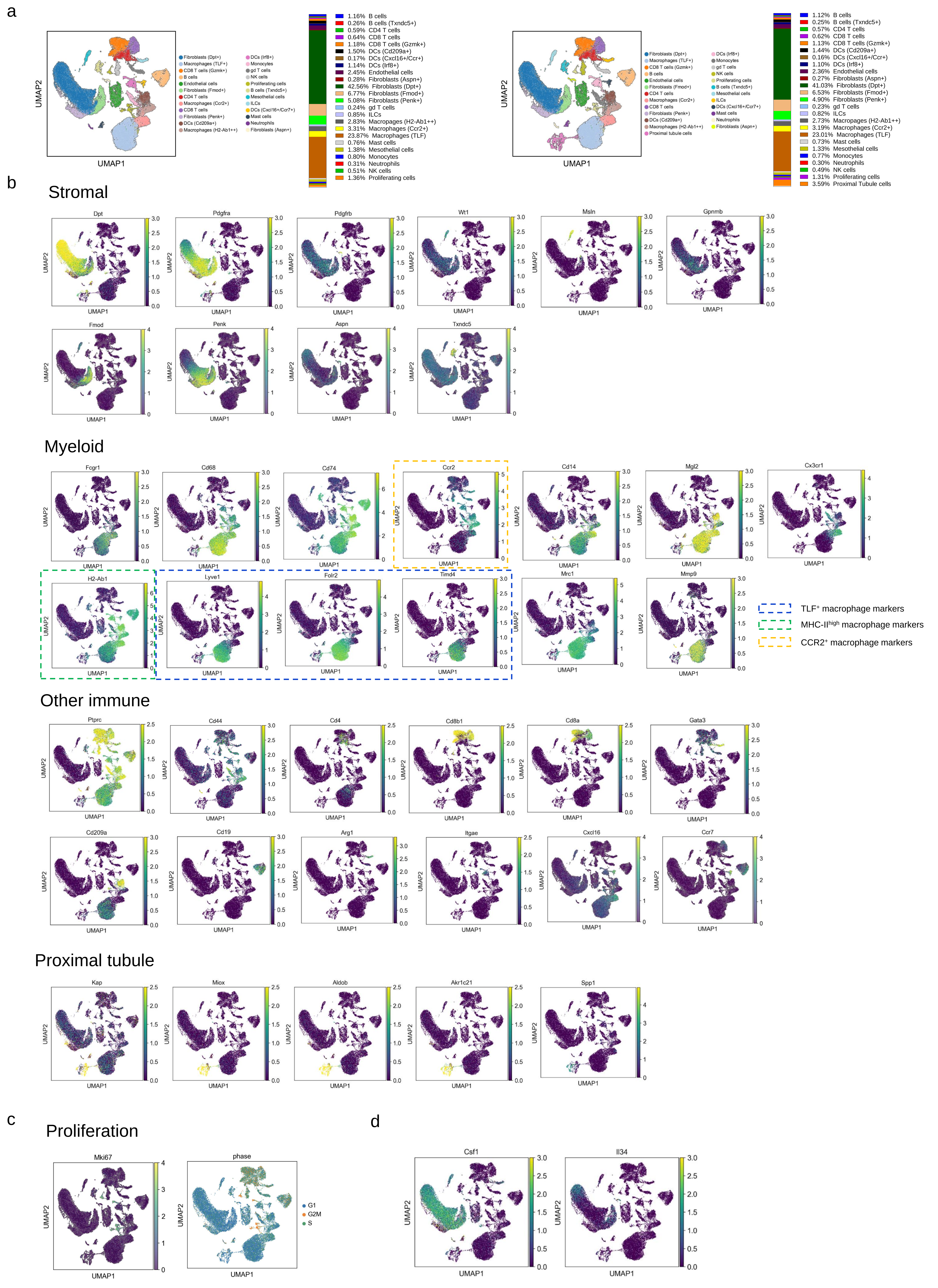

Supplementary Figure 6: Supporting bulk RNA-Seq and flow cytometry data for RCAMs and TLF$^{+}$ macrophages.

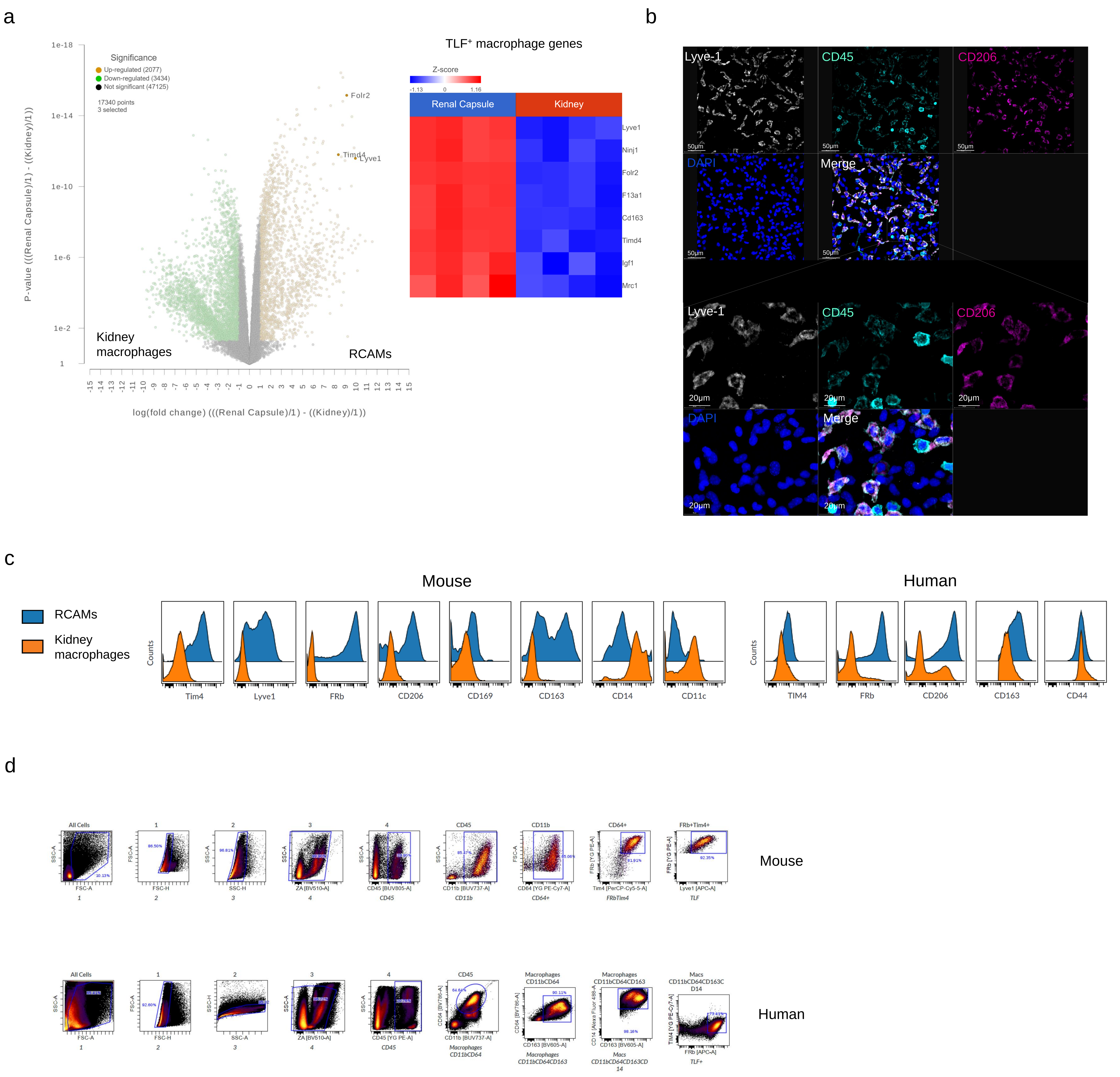

Supplementary Figure 7: RCAMs MHC-II expression in various conditions.

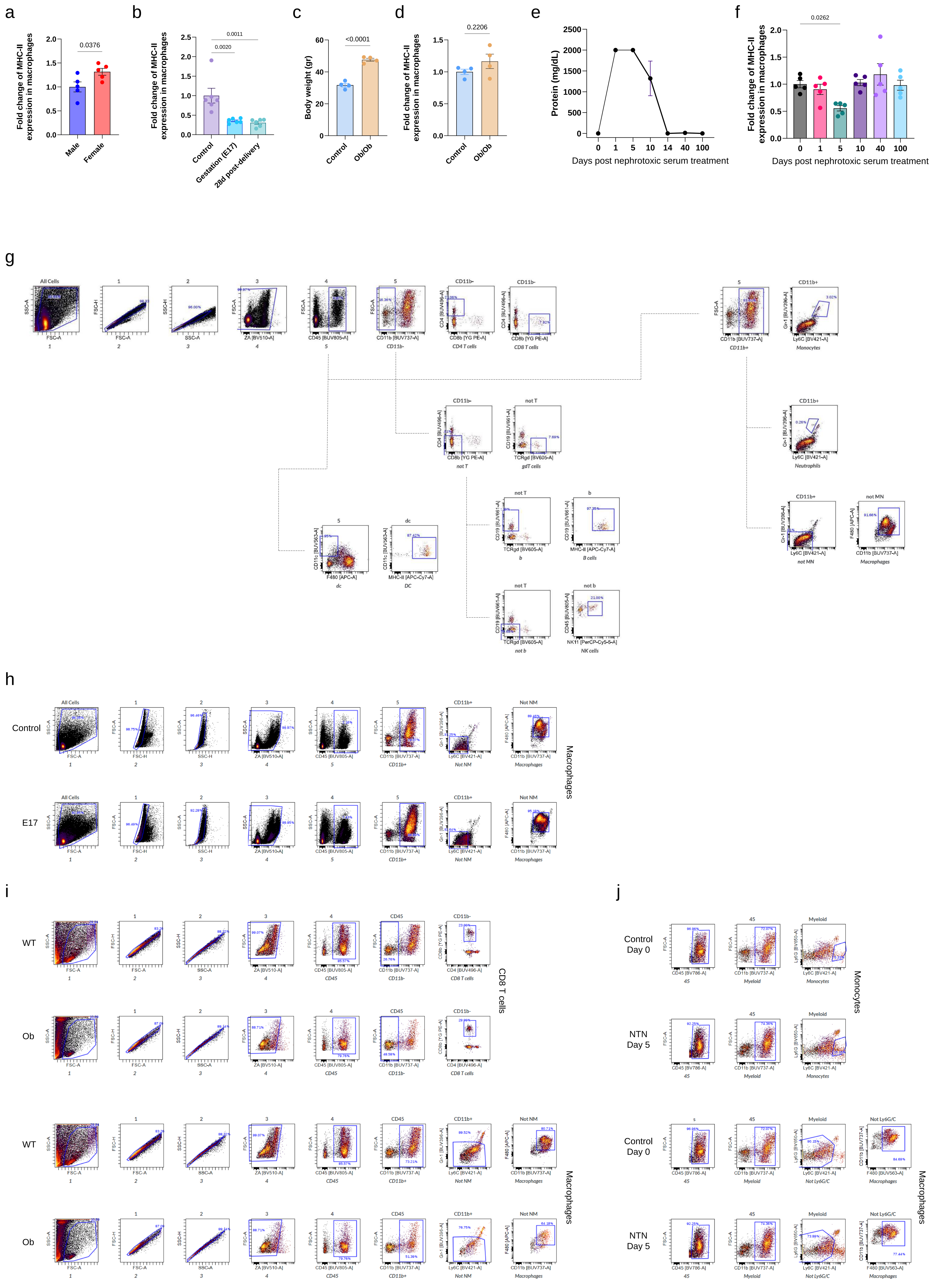

Supplementary Figure 8: Supporting histology, flow cytometry, and bulk RNA-Seq data of age-related changes in the renal capsule.

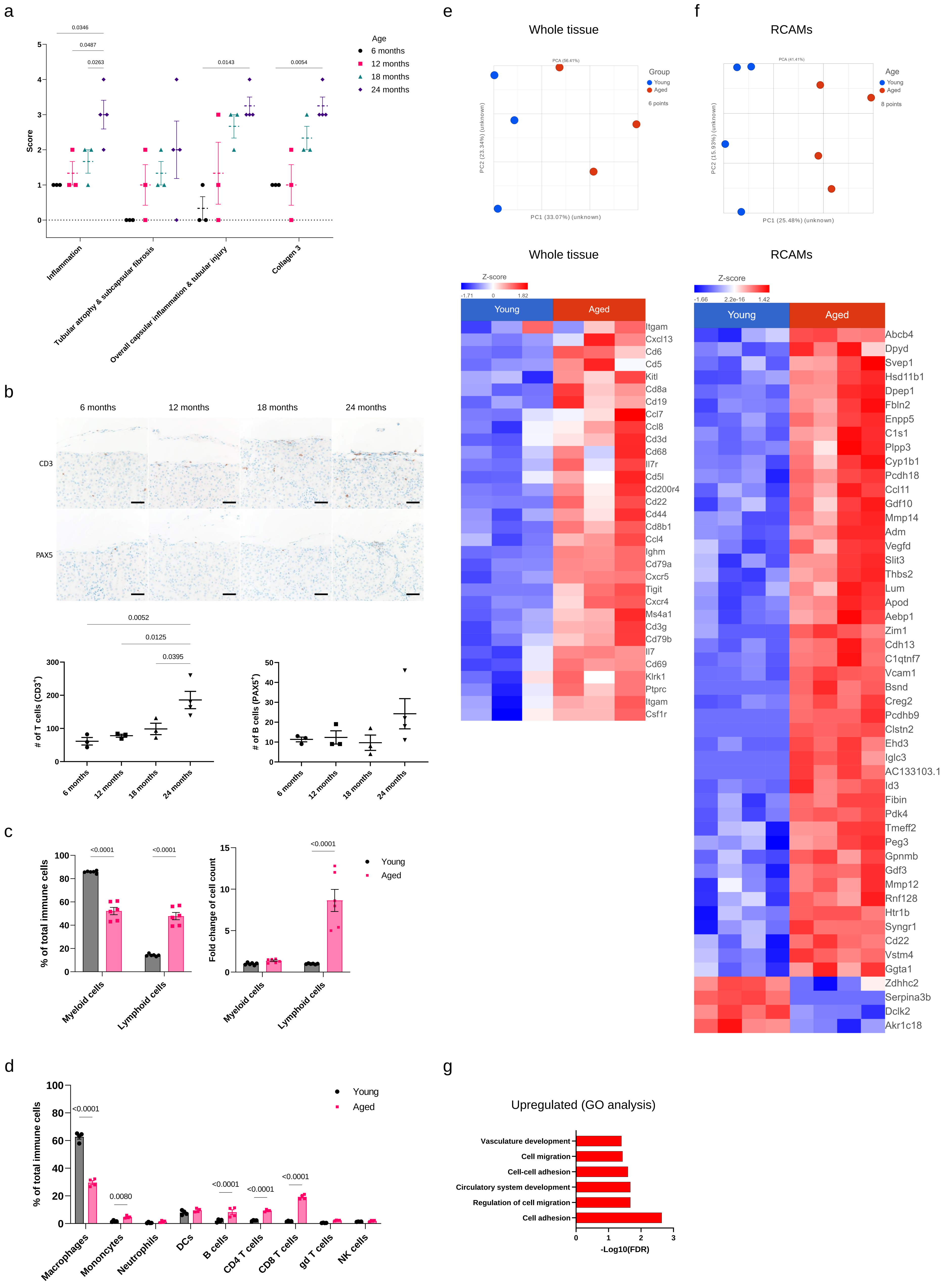

Supplementary Figure 9: Supporting data of age-related changes in the renal capsule.

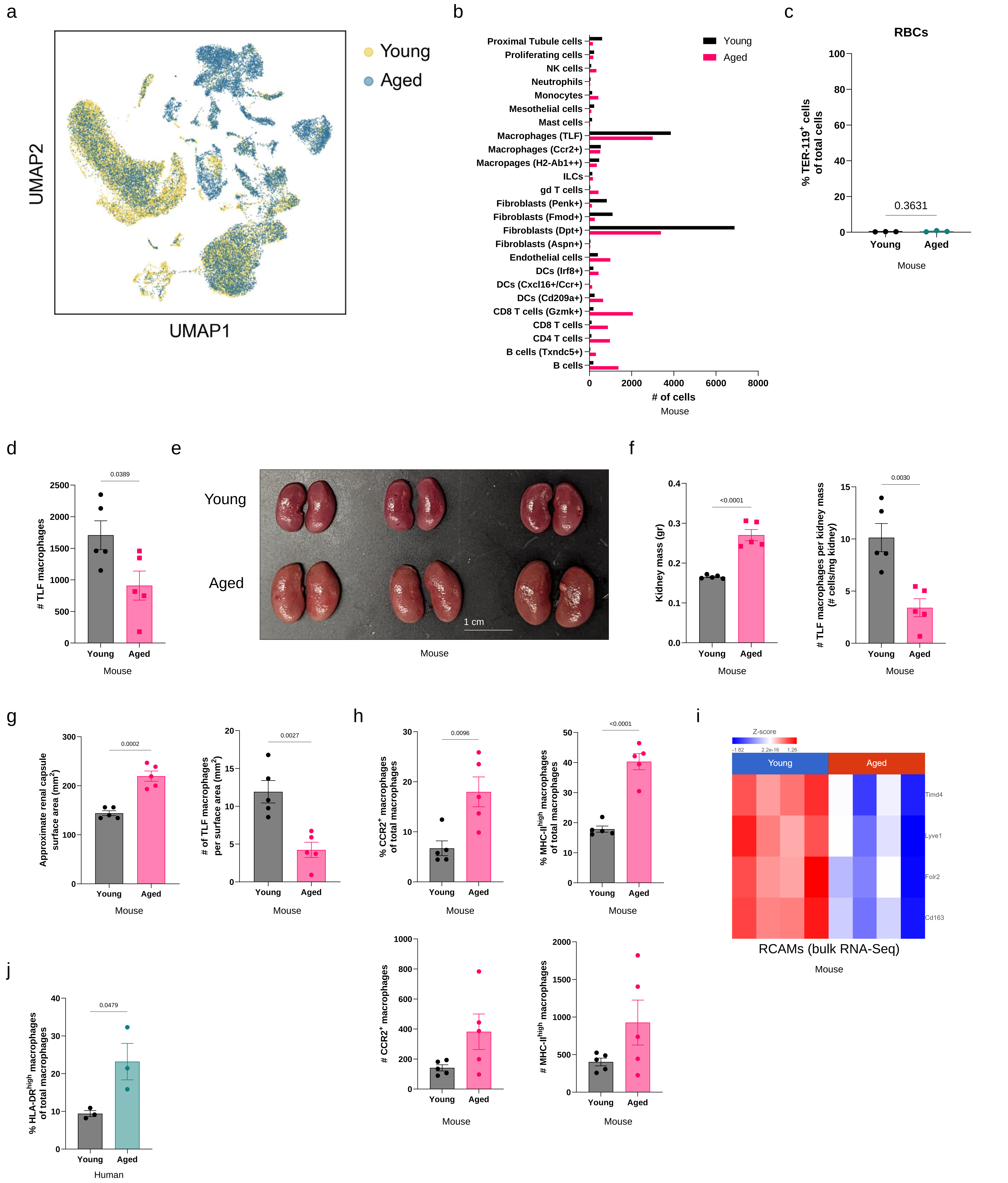

Supplementary Figure 10: Supporting data of age-related changes in the renal capsule, scRNA-Seq data.

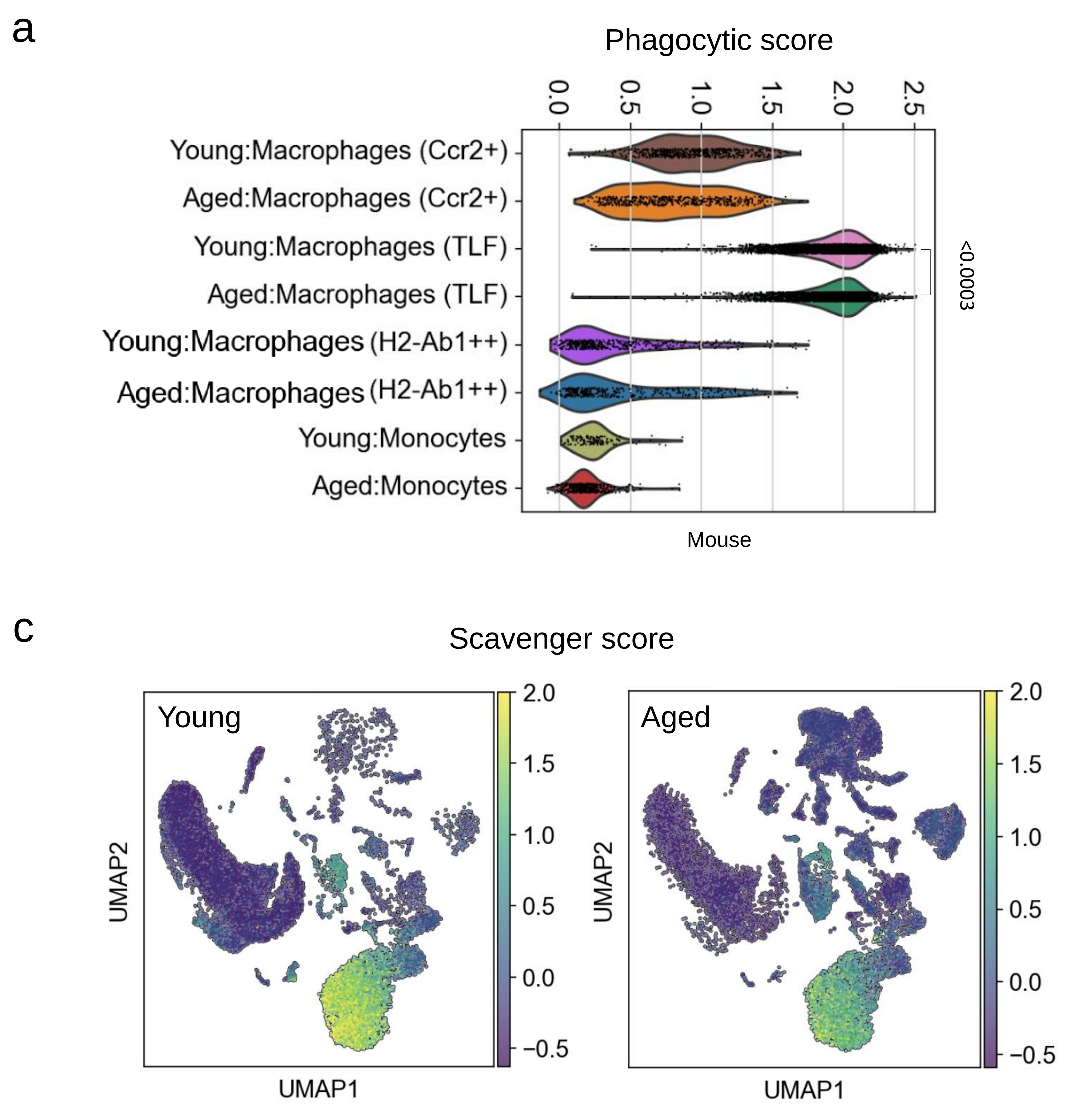

Supplementary Figure 11: Supporting data of 'aging score' related changes in the renal capsule.

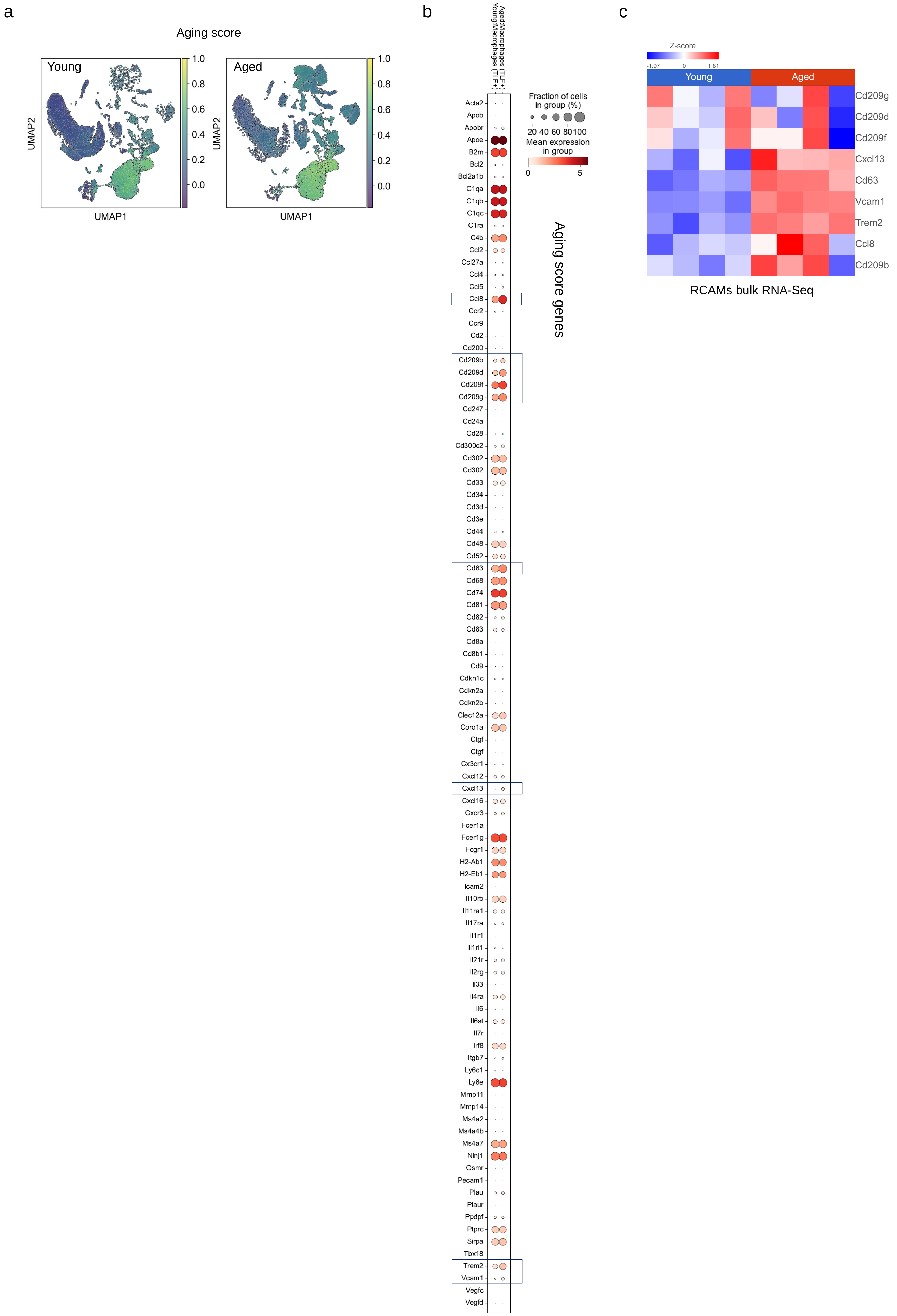

Supplementary Figure 12: Supporting data of SenMayo score related changes in the renal capsule.

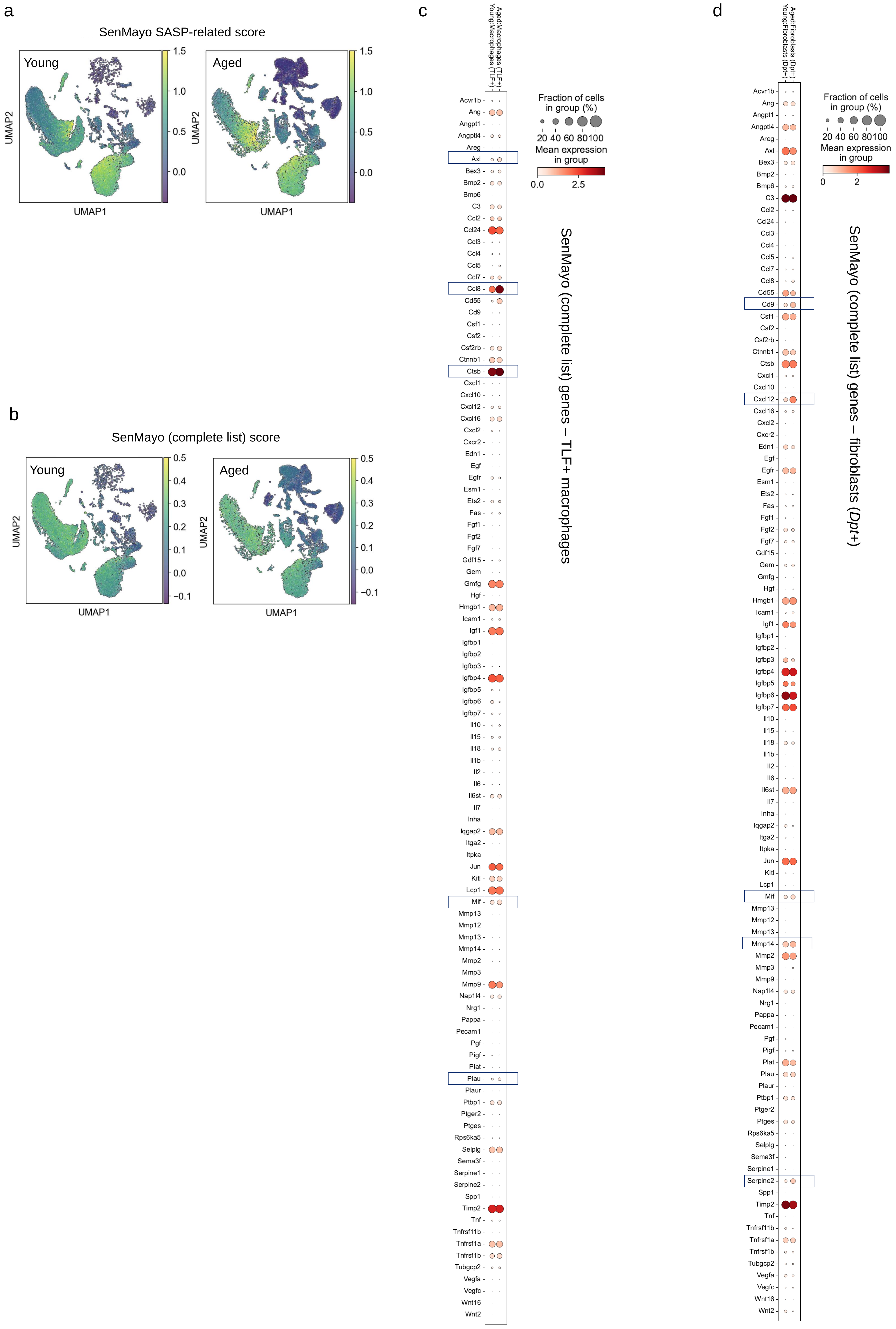

Supplementary Figure 13: Supporting data of 'SASP score' related changes in the renal capsule.

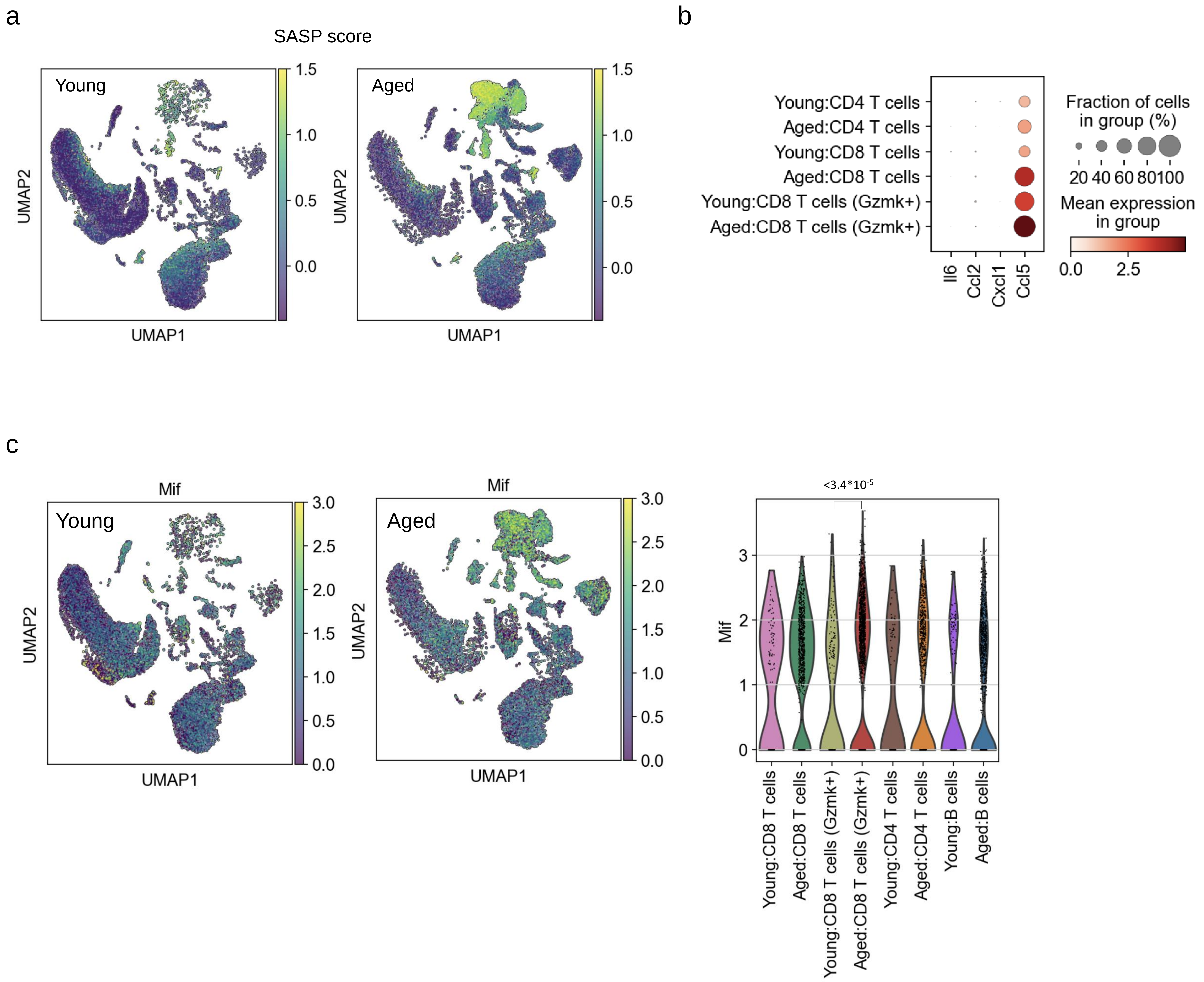

Supplementary Figure 14: Supporting data of renal capsule CellChat analysis.

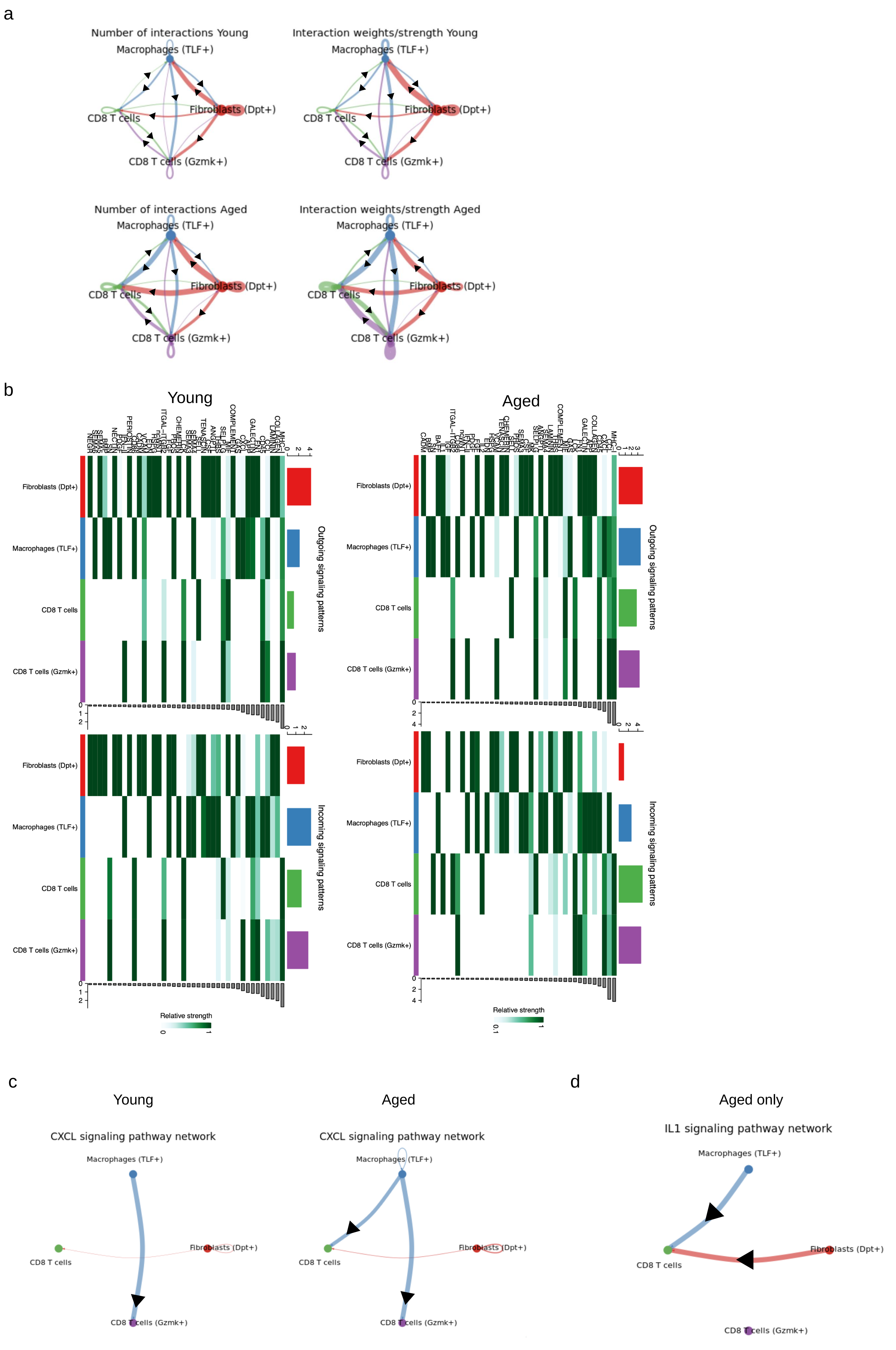

Supplementary Figure 15: Schematic description of suggested cell interactions in the renal capsule in aging.

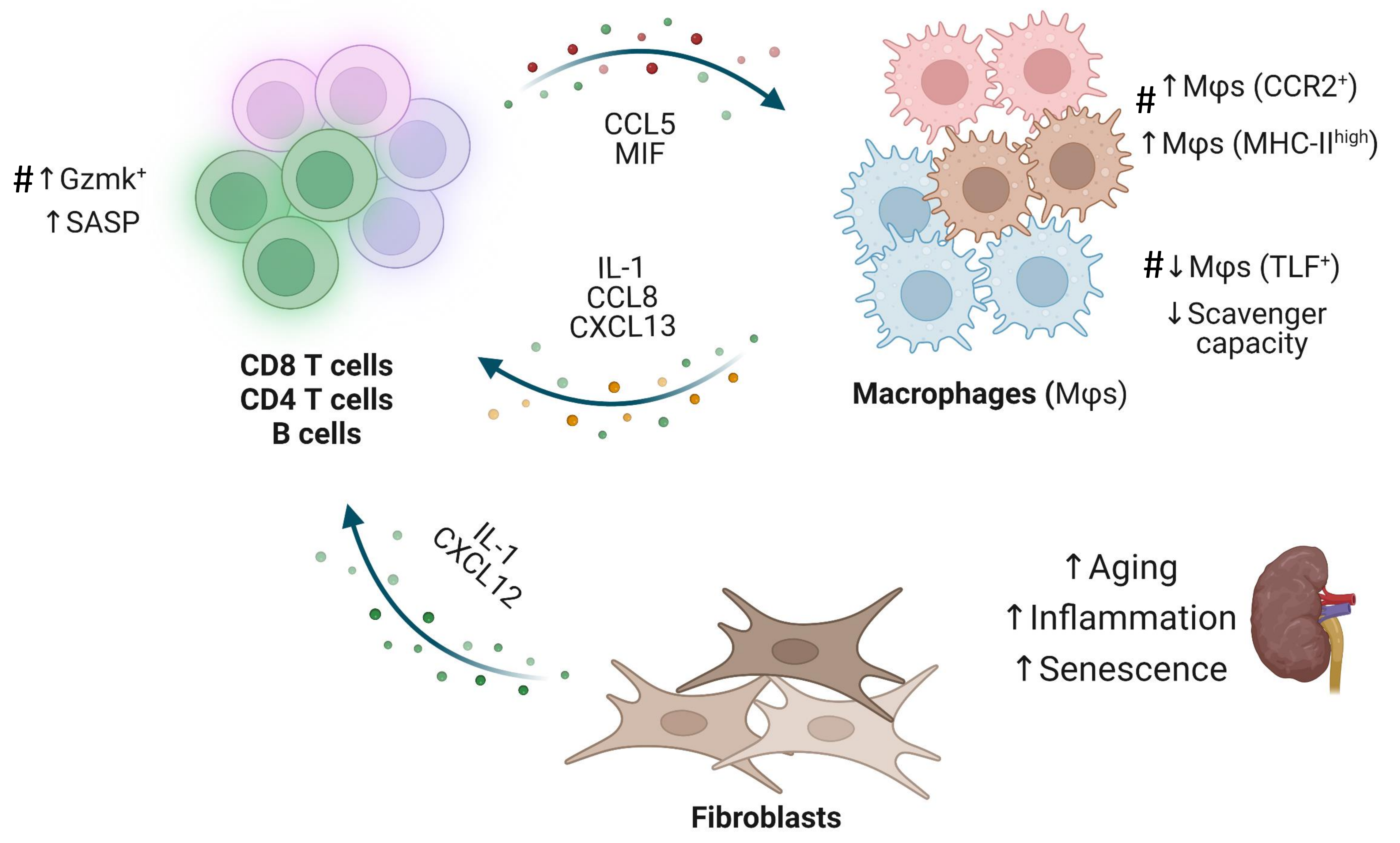
