## Supplementary Tables for "The renal capsule, a vibrant and adaptive cell environment of the kidney in homeostasis and aging"

**Supplementary Table 1**: Detailed information of flow cytometry markers used in this study.

**Mouse**

| Fluorophore | Target | Manufacture | Catalogue # | Cell type expression |
|---|---|---|---|---|
| BUV496 | CCR2 | BD Biosciences | 750043 | CCR2+ macrophages |
| BUV661 | CCR2 | BD Biosciences | 750472 | CCR2+ macrophages |
| BUV737 | CD11b | BD Biosciences | 741722 | Myeloid cells |
| Per-CP Cy5.5 | CD11b | Biolegend | 101228 | Myeloid cells |
| BUV563 | CD11c | BD Biosciences | 749040 | Dendritic cells (DCs) |
| BUV563 | CD11c | BD Biosciences | 749040 | Dendritic cells (DCs) |
| PE-Cy7 | CD11c | Biolegend | 117318 | Dendritic cells (DCs) |
| PE | CD14 | Biolegend | 123310 | Myeloid cells |
| PE-Cy7 | CD140a | Biolegend | 135912 | Fibroblasts |
| BV421 | CD163 | Biolegend | 155309 | TLF+ macrophages |
| APC | CD163 | Biolegend | 155306 | TLF+ macrophages |
| BV605 | CD169 | Biolegend | 142413 | Macrophages |
| BUV661 | CD19 | BD Biosciences | 612971 | B cells |
| BV711 | CD206 | Biolegend | 141727 | TLF+ macrophages |
| AF488 | CD206 | Biolegend | 141710 | TLF+ macrophages |
| BV650 | CD3 | Biolegend | 100229 | T cells |
| AF488 | CD31 | Biolegend | 102414 | Endothelial cells |
| BUV496 | CD4 | BD Biosciences | 741051 | CD4 T cells |
| BUV805 | CD45 | BD Biosciences | 752415 | Leukocytes |
| APC | CD45 | Biolegend | 103112 | Leukocytes |
| APC-Cy7 | CD45 | Biolegend | 103116 | Leukocytes |
| BV786 | CD64 | BD Biosciences | 741024 | Macrophages |
| PE-Cy7 | CD64 | Biolegend | 139314 | Macrophages |
| PE | CD8b | Biolegend | 126608 | CD8 T cells |
| AF488 | CX3CR1 | Biolegend | 149022 | Macrophages |
| APC | F4/80 | Biolegend | 123116 | Macrophages |
| BUV563 | F4/80 | BD Biosciences | 749284 | Macrophages |
| BUV396 | F480 | BD Biosciences | 565614 | Macrophages |
| BUV496 | F480 | BD Biosciences | 750644 | Macrophages |
| PE | FRb | Biolegend | 153304 | TLF+ macrophages |
| BUV395 | Gr-1 | BD Biosciences | 563849 | Neutrophils |
| Zombie Aqua | Live/dead | Biolegend | 423102 | Live/Dead cells |
| BV421 | Ly6C | Biolegend | 128032 | Monocytes |
| AF488 | Ly6C | Biolegend | 128022 | Monocytes |
| BV605 | Ly6G | Biolegend | 127639 | Neutrophils |
| BV650 | Ly6G | Biolegend | 127641 | Neutrophils |
| eFluor 660 | Lyve-1 | Invitrogen | 50-0443-82 | TLF+ macrophages |
| APC-Cy7 | MHC-II | Biolegend | 117324 | MHC-II High macrophages |
| PerCP-Cy5.5 | NK1.1 | Biolegend | 156526 | NK cells |
| BUV805 | TCRb | BD Biosciences | 748405 | T cells |
| BV605 | TCRgd | Biolegend | 118129 | gd T cells |
| BV650 | TER-119 | Biolegend | 116235 | Erythrocytes |
| PerCP-Cy5.5 | Tim4 | Biolegend | 130020 | TLF+ macrophages |
| BUV737 | Tim4 | BD Biosciences | 749134 | TLF+ macrophages |

**Human**

| Fluorophore | Target | Manufacture | Catalogue # | Cell type expression |
|---|---|---|---|---|
| APC-Cy7 | CCR2 | Biolegend | 357220 | CCR2+ macrophages |
| BUV737 | CD11b | BD Biosciences | 741826 | Myeloid cells |
| BV605 | CD11c | Biolegend | 301636 | Dendritic cells (DCs) |
| AF488 | CD14 | Biolegend | 301811 | Myeloid cells / Monocytes |
| PE-Cy7 | CD15 | Biolegend | 323030 | Neutrophils |
| BV605 | CD163 | Biolegend | 333616 | Macrophages |
| BUV396 | CD19 | BD Biosciences | 563549 | B cells |
| APC | CD206 | Biolegend | 321110 | Macrophages |
| PerCP-Cy5.5 | CD3 | Biolegend | 344808 | T cells |
| BUV496 | CD4 | BD Biosciences | 741134 | CD4 T cells |
| APC-Cy7 | CD44 | Biolegend | 103028 | Leukocytes |
| PE | CD45 | Biolegend | 304008 | Leukocytes |
| BUV563 | CD56 | BD Biosciences | 612929 | NK cells |
| BV786 | CD64 | Biolegend | 305044 | Macrophages |
| BUV805 | CD8 | BD Biosciences | 612889 | CD8 T cells |
| APC | FRb | Biolegend | 391706 | TLF+ macrophages |
| BUV661 | HLA-DR | BD Biosciences | 612981 | HLA-DR High macrophages |
| Zombie Aqua | Live/dead | Biolegend | 423102 | Live/Dead cells |
| BV711 | TCRab | Biolegend | 306740 | T cells |
| BV421 | TCRgd | Biolegend | 331218 | gd T cells |
| PE-Cy7 | TIM4 | Biolegend | 354006 | TLF+ macrophages |

**Supplementary Table 2**: Detailed information of mass cytometry markers used in this study.

| Metal isotope | Target | Manufacture | Catalogue # | Surface/intracellular staining |
|---|---|---|---|---|
| 103Rh | Live/dead | Fluidigm | 201103B | NA |
| 141Pr | Ly6G | Fluidigm | 3141008B | Surface |
| 142Nd | CXCR5 | Fluidigm | 3142015B | Surface |
| 143Nd | TCRb | Fluidigm | 3143010B | Surface |
| 144Nd | MHC-I | Fluidigm | 3144016B | Surface |
| 145Nd | CD69 | Fluidigm | 3145005B | Surface |
| 146Nd | F4/80 | Fluidigm | 3146008B | Surface |
| 147Sm | CD45 | Fluidigm | 3147003B | Surface |
| 148Nd | CD140a | Fluidigm | 3148018B | Surface |
| 149Sm | CD19 | Fluidigm | 3149002B | Surface |
| 150Nd | Ly6C | Fluidigm | 3150010B | Surface |
| 151Eu | CD64 | Fluidigm | 3151012B | Surface |
| 152Sm | CD3e | Fluidigm | 3152004B | Surface |
| 153Eu | CD16/32 | Fluidigm | 3153011B | Surface |
| 154Sm | CD11b | Fluidigm | 3154006B | Surface |
| 156Gd | CD14 | Fluidigm | 3156009B | Surface |
| 158Gd | IL-10 | Fluidigm | 3158002B | Intracellular |
| 159Tb | CXCR4 | Fluidigm | 3159030B | Surface |
| 160Gd | CD5 | Fluidigm | 3160002B | Surface |
| 161Dy | CD40 | Fluidigm | 3161020B | Surface |
| 162Dy | Tim3 | Fluidigm | 3162029B | Surface |
| 163Dy | ICAM-1 (CD54) | Fluidigm | 3163020B | Surface |
| 164Dy | CX3CR1 | Fluidigm | 3164023B | Surface |
| 165Ho | PECAM-1 (CD31) | Fluidigm | 3165013B | Surface |
| 166Er | EpCAM (CD326) | Fluidigm | 3166014B | Surface |
| 167Er | IL-6 | Fluidigm | 3167003B | Intracellular |
| 168Er | CD8a | Fluidigm | 3168003B | Surface |
| 169Tm | CD206 | Fluidigm | 3169021B | Surface |
| 170Er | CD40L | Fluidigm | 3170011B | Surface |
| 171Yb | CD44 | Fluidigm | 3171003B | Surface |
| 172Yb | CD4 | Fluidigm | 3172003B | Surface |
| 173Yb | c-kit | Fluidigm | 3173004B | Surface |
| 174Yb | MHC-II | Fluidigm | 3174003B | Surface |
| 175Lu | CD38 | Fluidigm | 3175014B | Surface |
| 176Lu | B220 | Fluidigm | 3176002B | Surface |
| 191/193Ir | Live/dead | Fluidigm | 201192B | NA |
| 209Bi | CD11c | Fluidigm | 3209005B | Surface |

**Supplementary Table 3**: Gene lists for scRNA-Seq scores described in this study.

| Inflammatory score | Phagocytic score | Scavenger score | Aging score | SenMayo SASP-related score | SenMayo Complete list score | SASP score |
|---|---|---|---|---|---|---|
| *Msrb1* | *Fcna* | *Cd163* | *Acta2* | *Ccl2* | *Acvr1b* | *Il6* |
| *Hp* | *Folr2* | *Cd38* | *Apob* | *Cxcl14* | *Ang* | *Ccl2* |
| *Gm9733* | *Lyve1* | *Lyve1* | *Apobr* | *Cxcl12* | *Angpt1* | *Cxcl1* |
| *Samhd1* | *Cd209f* | *Msr1* | *Apoe* | *Hp* | *Angptl4* | *Ccl5* |
| *Prdx5* | *Fxyd2* | *Cd36* | *B2m* | *Trf* | *Areg* | |
| *Plac8* | *Cd36* | | *Bcl2* | *Serping1* | *Axl* | |
| *Esd* | *Ltc4s* | | *Bcl2a1b* | *Mt1* | *Bex3* | |
| *Filip1l* | *Ednrb* | | *C1qa* | *Tmem176b* | *Bmp2* | |
| *Ifitm3* | *Sepp1* | | *C1qb* | *Mt2* | *Bmp6* | |
| *Nadk* | *Rcn3* | | *C1qc* | *Igfbp4* | *C3* | |
| *1600014C10Rik* | *Fcgrt* | | *C1ra* | *Grem1* | *Ccl1* | |
| *Lgals3* | *Cd209g* | | *C4b* | *Cd302* | *Ccl2* | |
| *Mcemp1* | *Wfdc17* | | *Ccl16* | *Apoe* | *Ccl20* | |
| *Daxx* | *Timd4* | | *Ccl2* | *Msmp* | *Ccl24* | |
| *Itgal* | *Ninj1* | | *Ccl27a* | *Adipoq* | *Ccl26* | |
| *Cd300a* | *Cbr2* | | *Ccl4* | *Cyr61* | *Ccl3* | |
| *Sell* | *Cd163* | | *Ccl5* | *Gas6* | *Ccl4* | |
| *Gm7676* | *Ccl24* | | *Ccl8* | *Mmp13* | *Ccl5* | |
| *Ms4a4c* | *F13a1* | | *Ccr2* | *Tmem176a* | *Ccl7* | |
| *Arpc1b* | *Mtss1* | | *Ccr9* | | *Ccl8* | |
| *Ms4a6c* | *Cd209d* | | *Cd2* | | *Cd55* | |
| *Pydc4* | *Clec10a* | | *Cd200* | | *Cd9* | |
| *Nfe2* | *Gas6* | | *Cd209b* | | *Csf1* | |
| *Aprt* | *C1qa* | | *Cd209d* | | *Csf2* | |
| *Pnp* | *Ctsb* | | *Cd209f* | | *Csf2rb* | |
| *Clec4e* | *Cfh* | | *Cd209g* | | *Cst10* | |
| *Fyb* | *C4b* | | *Clec12a* | | *Ctnnb1* | |
| *Psmb9* | *Abca1* | | *Coro1a* | | *Ctsb* | |
| *Ifitm6* | *Igfbp4* | | *Ctgf* | | *Cxcl1* | |
| *2310001H17Rik* | *Serpinb6a* | | *Ctgf* | | *Cxcl10* | |
| *B4galnt1* | *Mrc1* | | *Cx3cr1* | | *Cxcl12* | |
| *Taldo1* | *Timp2* | | *Cxcl12* | | *Cxcl16* | |
| *Agpat4* | *Blvrb* | | *Cxcl13* | | *Cxcl2* | |
| *Gngt2* | *Nrp1* | | *Cxcl16* | | *Cxcl3* | |
| *Fgr* | *Cmah* | | *Cxcr3* | | *Cxcr2* | |
| *Spi1* | *Csf1r* | | *Fcer1a* | | *Dkk1* | |
| *Chil3* | *C1qc* | | *Fcer1g* | | *Edn1* | |
| *Klf3* | *Aldh2* | | *Fcgr1* | | *Egf* | |
| *Itga4* | *Itm2b* | | *H2-Ab1* | | *Egfr* | |
| *Surf1* | *Wwp1* | | *H2-Eb1* | | *Ereg* | |
| *Tfec* | *Cd63* | | *Icam2* | | *Esm1* | |
| *Smpdl3a* | *AI607873* | | *Il10rb* | | *Ets2* | |
| *Tnfaip8l2* | *Emp1* | | *Il11ra1* | | *Fas* | |
| *Pyhin1* | *Serinc3* | | *Il17ra* | | *Fgf1* | |
| *Pglyrp1* | *Pf4* | | *Il1r1* | | *Fgf2* | |
| *Mpeg1* | *Snx2* | | *Il1rl1* | | *Fgf7* | |
| *Ifi35* | *Apoe* | | *Il21r* | | *Gdf15* | |
| *Pxk* | *Dab2* | | *Il2rg* | | *Gem* | |
| *Oas3* | *Tmem106a* | | *Il33* | | *Gmfg* | |
| *Pkm* | *Ophn1* | | *Il4ra* | | *Hgf* | |
| *Eml4* | *Pltp* | | *Il6* | | *Hmgb1* | |
| *Phkg2* | *Pmp22* | | *Il6r* | | *Icam1* | |
| *AI413582* | *Lgmn* | | *Cd247* | | *Icam5* | |
| *Fcgr1* | *Maf* | | *Cd24a* | | *Igf1* | |
| *Clec4d* | *Mt1* | | *Cd28* | | *Igfbp1* | |
| *Ifngr1* | *Ubc* | | *Cd300c2* | | *Igfbp2* | |
| *Anxa1* | *Cst3* | | *Cd302* | | *Igfbp3* | |
| *Metrnl* | *Pde2a* | | *Cd302* | | *Igfbp4* | |
| *Tyrobp* | *Prune2* | | *Cd33* | | *Igfbp5* | |
| *Tgm2* | *Bmp2* | | *Cd34* | | *Igfbp6* | |
| *Snrnp40* | *Trf* | | *Cd3d* | | *Igfbp7* | |
| *Dok3* | *Cltc* | | *Cd3e* | | *Il10* | |
| *Stk38* | *Glul* | | *Cd44* | | *Il13* | |
| *Ifi204* | *Ly6e* | | *Cd48* | | *Il15* | |
| *Slc16a3* | *Rgl1* | | *Cd52* | | *Il18* | |
| *Tpgs1* | *C5ar1* | | *Cd63* | | *Il1a* | |
| *AF251705* | *Pepd* | | *Cd68* | | *Il1b* | |
| *Lyz2* | *mt-Atp6* | | *Cd74* | | *Il2* | |
| *Tmem51* | *Stab1* | | *Cd81* | | *Il6* | |
| *Card19* | *Tgfbr2* | | *Cd82* | | *Il6st* | |
| *Slc33a1* | | | *Cd83* | | *Il7* | |
| *Isg20* | | | *Cd8a* | | *Inha* | |
| *Phf11d* | | | *Cd8b1* | | *Iqgap2* | |
| *Slpi* | | | *Cd9* | | *Itga2* | |
| *Hlcs* | | | *Cdkn1c* | | *Itpka* | |
| *Adrbk2* | | | *Cdkn2b* | | *Jun* | |
| *Gpr137b-ps* | | | *Il6st* | | *Kitl* | |
| *Gbp2* | | | *Il7r* | | *Lcp1* | |
| *Mrpl46* | | | *Irf8* | | *Mif* | |
| *Cytip* | | | *Itgb7* | | *Mmp13* | |
| *Rab8a* | | | *Ly6c1* | | *Mmp10* | |
| *Samsn1* | | | *Ly6e* | | *Mmp12* | |
| *Plgrkt* | | | *Mmp11* | | *Mmp13* | |
| *Oasl2* | | | *Mmp14* | | *Mmp14* | |
| *Lgals9* | | | *Ms4a2* | | *Mmp2* | |
| *Cdk2ap2* | | | *Ms4a4b* | | *Mmp3* | |
| *Ncor2* | | | *Ms4a7* | | *Mmp9* | |
| *Ldlrad3* | | | *Ninj1* | | *Nap1l4* | |
| *Sephs2* | | | *Osmr* | | *Nrg1* | |
| *Skint3* | | | *Pecam1* | | *Pappa* | |
| *Rnh1* | | | *Plau* | | *Pecam1* | |
| *I830077J02Rik* | | | *Plaur* | | *Pgf* | |
| *Fcgr3* | | | *Ppdpf* | | *Pigf* | |
| *B2m* | | | *Ptprc* | | *Plat* | |
| *Mndal* | | | *Sirpa* | | *Plau* | |
| *Ubxn2b* | | | *Tbx18* | | *Plaur* | |
| *Sema4d* | | | *Trem2* | | *Ptbp1* | |
| *Fnbp4* | | | *Vcam1* | | *Ptger2* | |
| *Rnase6* | | | *Vegfc* | | *Ptges* | |
| *Rrad* | | | *Vegfd* | | *Rps6ka5* | |
| *Glipr1* | | | | | *Scamp4* | |
| *Krr1* | | | | | *Selplg* | |
| *Rnf115* | | | | | *Sema3f* | |
| *Nuak2* | | | | | *Serpinb3a* | |
| *Ngdn* | | | | | *Serpine1* | |
| *Gm26740* | | | | | *Serpine2* | |
| *Thbs1* | | | | | *Spp1* | |
| *Coro1a* | | | | | *Spx* | |
| *Nsfl1c* | | | | | *Timp2* | |
| *Ethe1* | | | | | *Tnf* | |
| *Zfyve9* | | | | | *Tnfrsf11b* | |
| *Pou2f2* | | | | | *Tnfrsf1a* | |
| *Zfand2b* | | | | | *Tnfrsf1b* | |
| *S100a11* | | | | | *Tubgcp2* | |
| *Pi16* | | | | | *Vegfa* | |
| *Gtf3a* | | | | | *Vegfc* | |
| *Zcchc10* | | | | | *Vgf* | |
| *Ube2l6* | | | | | *Wnt16* | |
| *Hmgb2* | | | | | *Wnt2* | |
| *Txn1* | | | | | | |
| *Glrx* | | | | | | |
| *Rsad2* | | | | | | |
| *Pla2g4a* | | | | | | |
| *Ncf4* | | | | | | |
| *Usp18* | | | | | | |
| *Mief1* | | | | | | |
| *St7l* | | | | | | |
| *Tmsb10* | | | | | | |
| *Milr1* | | | | | | |
| *Rmdn1* | | | | | | |
| *Ly6e* | | | | | | |
| *Cyfip2* | | | | | | |
| *H3f3b* | | | | | | |
| *Thap11* | | | | | | |
| *Glyr1* | | | | | | |
| *Pmm2* | | | | | | |
| *Gnas* | | | | | | |
| *Tifab* | | | | | | |
| *Rap2b* | | | | | | |
| *Taf5* | | | | | | |
| *Fam63a* | | | | | | |
| *Pkn1* | | | | | | |
| *Zfp800* | | | | | | |
| *Atrn* | | | | | | |
| *Msr1* | | | | | | |
| *Lst1* | | | | | | |
| *Mxd1* | | | | | | |
| *Fabp5* | | | | | | |
| *Ripk3* | | | | | | |
| *Cyba* | | | | | | |
| *Pdzd11* | | | | | | |
| *Ogfr* | | | | | | |
| *Lilra6* | | | | | | |
| *Srgn* | | | | | | |
| *Pfn1* | | | | | | |
| *Psmb8* | | | | | | |
| *Zfp653* | | | | | | |
| *Aldh16a1* | | | | | | |
| *Med4* | | | | | | |
| *Prpf38b* | | | | | | |
| *Rilpl2* | | | | | | |
| *Egln2* | | | | | | |
| *Rnf138* | | | | | | |
| *Lmo4* | | | | | | |

**Supplementary Table 4**: Details regarding histology scores described in this study.

| | Inflammation |
|---|---|
| **0** | No significant inflammatory infiltrates |
| **1** | 1-2 small leukocyte aggregates (less than approximately 10 cells or scattered leukocytes associated with tubular atrophy) |
| **2** | 3-5 small leukocyte aggregates or 1 larger lymphocyte aggregate |
| **3** | 6-10 small leukocyte aggregates or 2-3 larger aggregates |
| **4** | >10 small leukocyte aggregates or >3 larger aggregates |
| **5** | Locally extensive inflammation |

| | Tubular Atrophy and subcapsular fibrosis |
|---|---|
| **0** | No subcapsular tubular atrophy |
| **1** | 1-2 foci of subcapsular tubular atrophy |
| **2** | 3-5 foci of subcapsular tubular atrophy |
| **3** | 6-10 foci of subcapsular tubular atrophy |
| **4** | >10 foci of subcapsular tubular atrophy |
| **5** | Locally tubular atrophy |

| | Overall Capsular Inflammation and Tubular Injury Score |
|---|---|
| **0** | No capsule-associated lesions observed |
| **1** | Minimal, focal capsule fibrosis or inflammation with capsule retraction |
| **2** | Focal cluster of atrophied tubules |
| **3** | Multifocal tubular atrophy and inflammation |
| **4** | Confluent regions of tubular atrophy and inflammation |

| | Collagen 3 |
|---|---|
| **0** | No abnormal thickening |
| **1** | 1-2 fibrotic foci |
| **2** | 3-5 fibrotic foci |
| **3** | > 5 fibrotic foci or confluent foci |
| **4** | extensive capsular fibrosis and remodeling |

| CCR2+ | | |
|---|---|---|
| names | [illegible] | pvals_adj |
| [illegible] | [illegible] | [illegible] |
| Gngt2 | 1.1784319 | 1.73E-08 |
| H2-Aa | 1.1715126 | 1.75E-47 |
| [illegible] | [illegible] | [illegible] |
| Tmem205 | 1.0297146 | 1.26E-03 |
| Ms4a7 | 0.9719795 | 2.54E-11 |
| [illegible] | [illegible] | [illegible] |
| Bcl2a1b | 0.60963256 | 1.20E-03 |
| [illegible] | 0.59735405 | 3.28E-05 |
| [illegible] | [illegible] | [illegible] |
| [illegible] | 0.25436693 | 1.72E-03 |

| MHC-II high | | |
|---|---|---|
| names | [illegible] | pvals_adj |
| [illegible] | [illegible] | [illegible] |
| Fcgr1 | 1.9746988 | 2.32E-04 |
| Cd63 | 1.9150531 | 2.22E-13 |
| [illegible] | [illegible] | [illegible] |
| Cd300c2 | 1.7298508 | 2.69E-21 |
| Ccl8 | 1.7154235 | 1.00E-03 |
| [illegible] | [illegible] | [illegible] |
| Rgs1 | 1.2121505 | 5.28E-05 |
| Grk3 | 1.1927134 | 4.97E-03 |
| [illegible] | [illegible] | [illegible] |
| Il18 | 0.9168537 | 2.03E-03 |
| [illegible] | [illegible] | [illegible] |
| Eif1 | 0.25204492 | 4.94E-06 |

| TLF+ | | |
|---|---|---|
| names | [illegible] | pvals_adj |
| [illegible] | [illegible] | [illegible] |
| Stx3 | 3.1926818 | 3.98E-06 |
| Ccl8 | 3.118745 | 0.00E+00 |
| [illegible] | [illegible] | [illegible] |
| Gm7609 | 2.9172268 | 5.45E-12 |
| Pcdhac2 | 2.8751655 | 1.74E-04 |
| [illegible] | [illegible] | [illegible] |
| Vsig4 | 1.9393672 | 1.72E-09 |
| Ndn | 1.9322537 | 3.37E-03 |
| [illegible] | [illegible] | [illegible] |
| Arsg | 1.248389 | 1.47E-22 |
| [illegible] | [illegible] | [illegible] |
| Bloc1s6 | 0.35813832 | 1.35E-03 |
| [illegible] | [illegible] | [illegible] |
| [illegible] | 0.25003403 | 5.83E-05 |

[illegible]

### CXCL

| interaction_name | pathway_name | ligand | receptor | agonist | antagonist | co_A_receptor | co_I_receptor | evidence | annotation | interaction_name_2 |
|---|---|---|---|---|---|---|---|---|---|---|
| [illegible] | [illegible] | [illegible] | [illegible] | [illegible] | [illegible] | [illegible] | [illegible] | [illegible] | [illegible] | [illegible] |

## IL1

| interaction_name | pathway_name | ligand | receptor | agonist | antagonist | co_A_receptor | co_I_receptor | evidence | annotation | interaction_name_2 |
|---|---|---|---|---|---|---|---|---|---|---|
| [illegible] | [illegible] | [illegible] | [illegible] | [illegible] | [illegible] | [illegible] | [illegible] | [illegible] | [illegible] | [illegible] |

### CCL

| interaction_name | pathway_name | ligand | receptor | agonist | antagonist | co_A_receptor | co_I_receptor | evidence | annotation | interaction_name_2 |
|---|---|---|---|---|---|---|---|---|---|---|
| [illegible] | [illegible] | [illegible] | [illegible] | [illegible] | [illegible] | [illegible] | [illegible] | [illegible] | [illegible] | [illegible] |

### MHC-I

| interaction_name | pathway_name | ligand | receptor | agonist | antagonist | co_A_receptor | co_I_receptor | evidence | annotation | interaction_name_2 |
|---|---|---|---|---|---|---|---|---|---|---|
| [illegible] | [illegible] | [illegible] | [illegible] | [illegible] | [illegible] | [illegible] | [illegible] | [illegible] | [illegible] | [illegible] |

### COLLAGEN

| interaction_name | pathway_name | ligand | receptor | agonist | antagonist | co_A_receptor | co_I_receptor | evidence | annotation | interaction_name_2 |
|---|---|---|---|---|---|---|---|---|---|---|
| [illegible] | [illegible] | [illegible] | [illegible] | [illegible] | [illegible] | [illegible] | [illegible] | [illegible] | [illegible] | [illegible] |

### LAMININ

| interaction_name | pathway_name | ligand | receptor | agonist | antagonist | co_A_receptor | co_I_receptor | evidence | annotation | interaction_name_2 |
|---|---|---|---|---|---|---|---|---|---|---|
| [illegible] | [illegible] | [illegible] | [illegible] | [illegible] | [illegible] | [illegible] | [illegible] | [illegible] | [illegible] | [illegible] |
